# Developmental bias explains the evolutionary trend towards simple leaf shapes

**DOI:** 10.1101/2025.08.17.670617

**Authors:** James S. Malone, Nora S. Martin, Samuel H. A. von der Dunk, Liliana M. Dávalos, Ard A. Louis

## Abstract

The relative influence of developmental bias and natural selection on evolutionary outcomes remains a central, unresolved question in evolutionary biology. Here, we combine large-scale phylogenetic trees with herbarium data to confirm that, across the evolutionary history of angiosperms, transitions from complex to simple leaf shapes have occurred at a significantly higher rate than the reverse. To investigate whether this directional trend reflects an underlying developmental bias, we simulate evolutionary transitions by introducing random mutations into a well-established computational model of leaf development. Despite excluding selection, the transition rates produced by the model closely align with those observed in the phylogenetic data. This agreement indicates that a strong developmental bias, reflecting the relative ease with which gene regulatory networks generate simple over complex leaves, has played a major role in shaping the evolution of leaf morphology. Our findings echo recent work showing that similar biases towards simplicity influence the evolution of molecular structures, including RNA secondary structures and protein quaternary assemblies. Together, these results suggest that a developmental bias favouring simplicity may be a widespread and general feature of evolutionary systems.

## I. INTRODUCTION

Much of the phenotypic diversity across the tree of life can be explained by adaptation to ecological demands via natural selection. However, this process is constrained by the mechanisms that generate phenotypic variation, including development [1, 2], which translates genetic information into form. While this idea is widely acknowledged in evolutionary developmental biology [1–3], only recently have advances in methodology and our understanding of development allowed researchers to examine its predictive power for evolutionary outcomes [4–11]. Yet it remains unclear how widespread and strong this developmental bias is across phylogeny [12, 13], particularly for complex traits like morphology. Here, we test whether developmental bias can account for macroevolutionary patterns in angiosperm leaf shape evolution.

Angiosperm leaves are an attractive system to study developmental bias. They display vast morphological diversity, much of which can be effectively represented in only two spatial dimensions due to minimal growth along the adaxial-abaxial (upper-lower surface) axis [14]. This diversity arises from variations in growth patterns regulated by gene regulatory networks (GRNs) that orchestrate both local and global growth dynamics during leaf development, allowing complex forms to emerge from anisotropic expansion. These processes include deeply conserved genes and developmental mechanisms [15]. Variation in leaf morphology is driven not so much by changes in gene content, but rather by differences in the spatial and temporal expression of shared developmental regulators. This helps explain why closely related species can display wide variation in leaf shape. For instance, the genus *Pelargonium* includes species with unlobed, lobed, dissected, and compound leaves [16]. Similarly, significant variation can occur within a single organism, as seen in *Morus alba* (mulberry), which produces both unlobed and lobed leaves, often with varying degrees of lobe symmetry [14, 17]. The conservation of basic growth mechanisms also suggests that some developmental biases will be, to first order, shared across many different angiosperm species, enabling largescale studies of the role of development in determining evolutionary outcomes.

At the same time, there is evidence that leaf shape can be influenced by ecological demands such as thermoregulation [18, 19], hydraulic efficiency [20], light interception [21–23] and herbivory [24, 25]. However, it is still unclear how strong and general these effects are in natural environments [26, 27]. Size and the presence/absence of teeth on the margin are often stronger predictors of ecology than the broader leaf dissection categories we use here [28–33] (see section S4 for further discussion).

In a notable study, Geeta *et al*. [39] analysed a phylogeny of 560 angiosperms, showing that most leaf shapes (76% in their sample—table II) are unlobed, and that transition rates between leaf shapes are highly asymmetric. Specifically, transitions from more complex forms—such as lobed, dissected, or compound leaves—to simpler unlobed forms occurred far more frequently than transitions in the opposite direction. What explains this global bias towards simpler leaves? Is it primarily development or is it primarily adaptation?

In order to answer this question, we proceed with two parallel lines of work. In the first, we extend the phylogenetic analysis of Geeta *et al*. [39] with two much larger, recent phylogenies [40, 41] and our own shape classification dataset made from 8,911 herbarium images [34]. These confirm that transitions to unlobed leaves are much more likely than transitions away from unlobed leaves.

In the second line of work, we examine the role of random mutations using a well-established computational model of angiosperm leaf development by Runions *et al*. [14] (fig. 5). This model captures key conserved developmental mechanisms thought to be broadly shared across angiosperms (Methods section IV A), allowing us to explore the generic effects of neutral evolution in the absence of direct selection on leaf shape. Remarkably, we find that random mutation alone is sufficient to reproduce the relative phylogenetic rates towards unlobed leaf forms. This suggests that developmental bias may be the first-order explanation of this macroevolutionary trend.

The basic mechanism driving this bias towards simplicity is straightforward [8]: complex leaf shapes need more tightly constrained GRNs than the simpler unlobed leaf shape [43]. As a result, random genetic variation is more likely to produce simple shapes rather than complex ones that require more finely tuned parameter combinations.

## II. RESULTS

### A. Transition rates from complex leaves to simple leaves are greater than the reverse across angiosperm phylogeny

To establish whether there is a phylogenetic bias towards simple, unlobed leaves, we inferred the rate of evolutionary transitions between different leaf shapes across angiosperm phylogeny. We first classified 8,911 herbarium images (fig. 1a) [34] into 4 discrete categories—unlobed, lobed, dissected and compound. We used these data, as well as another shape classification dataset from Geeta *et al*. [39], to label the tips of 5 phylogenetic trees representing angiosperm evolution [39–41]. Of these trees, 3 were resolved at the level of genus (Zuntini *et al*. [41] genus, Janssens *et al*. [40] genus and Geeta *et al*. [39]). The other two were resolved at the level of species (Zuntini *et al*. [41] species and Janssens *et al*. [40] species) enabling us to examine how altering the amount of species-level diversity affects inferred transition rates. In agreement with Geeta *et al*. [39], we found that the vast majority of the taxa represented at the tips of these new phylogenies had unlobed leaves (table II), already suggesting a strong bias towards simple, unlobed leaves. To infer the rates of evolution among these 4 categories while accounting for tree topology, we fitted a continuous-time Markov chain (CTMC) model to the labelled trees (Methods section IV E), using a Markov chain Monte Carlo (MCMC) approach implemented in BayesTraits V4.1.2 [42]. We detected significantly higher rates of transition to the simplest shape (unlobed) than away from it towards more complex shapes, as also found in Geeta *et al*. [39] (figs. 1a and 7a).

**FIG. 1.**
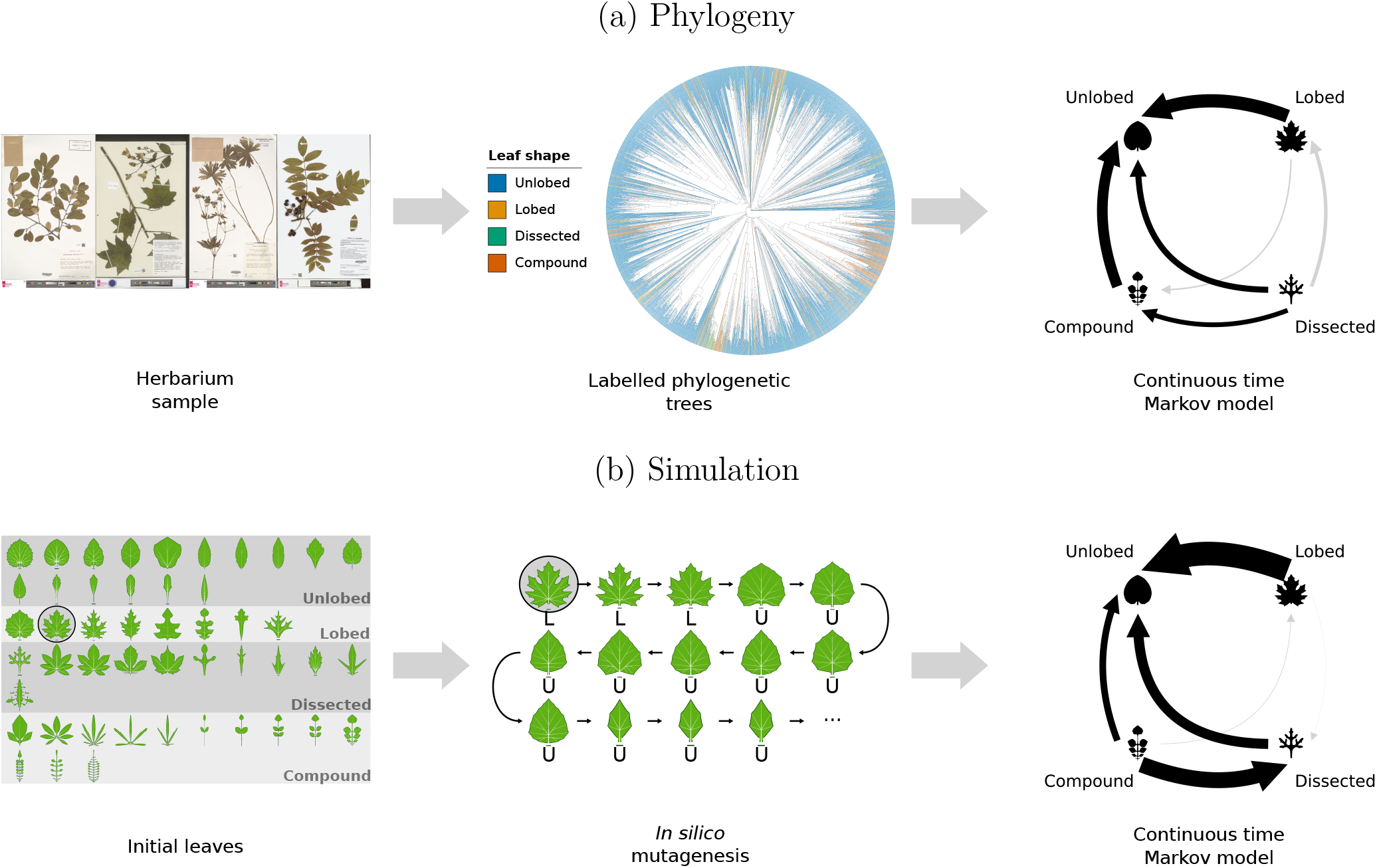
A summary of the methods and main results. We inferred transition rates between leaf shapes in two different ways: **(a)** by fitting a continuous-time Markov chain (CTMC) model of trait evolution to phylogenetic trees and **(b)** by simulating mutation in a computational model of leaf development. **(a)** (Left) Images sampled from the Naturalis Biodiversity Center herbarium [34–38] are manually classified as unlobed (u), lobed (l), dissected (d) or compound (c) (Methods section IV F). This dataset, as well as an analogous dataset by Geeta *et al*. [39], is used to label the tips of 5 different phylogenetic trees representing angiosperm evolution [39–41]. (Centre) The Zuntini *et al*. [41] species tree is shown in full. Rates are inferred by fitting a CTMC model to the labelled trees using a Markov chain Monte Carlo (MCMC) approach in BayesTraits V4.1.2 [42]. (Right) Posterior net rates for the Zuntini *et al*. [41] species tree are shown as arrows. Black (grey) arrows depict rates for which 0 falls outside (within) a 95% credible interval. **(b)** (Left) An initial sample of 48 leaves representing a wide morphospace is generated using the Runions *et al*. [14] model (Methods sections IV A and IV B). Leaves in this sample are mutated in parallel through a random walk algorithm, with no selection besides rejecting parameter combinations that give physically implausible leaves (table I), generating a sequence of leaves that evolve away from the initial leaf (Methods section IV C). (Middle) Every leaf is classified as unlobed (u), lobed (l), dissected (d) or compound (c) using an automated morphometric approach (Methods section IV D), giving a sequence of character states for each walk. To this trajectory, a CTMC model is fitted using MCMC (Methods section IV E). (Right) Arrows depict posterior net rates for the MUT2 scheme (alg. 1).

The bias towards unlobed shapes was consistent across the five phylogenetic trees (figs. 7a and 7b). We found no cases where the direction of two statistically significant (black arrows—where 0 falls outside a 95% credible interval) net rates disagreed between phylogenies (fig. 7a).

### B. computational model of leaf development recapitulates *in vivo* experiments demonstrating developmental bias towards unlobed leaf shape

To investigate how much of the preference for unlobed leaves can be explained by developmental bias, we use a popular model from Runions *et al*. [14] that combines gene-regulatory networks and key growth mechanisms for leaf shape (Methods section IV A). As a first validation test, we use the model to analyse *in vivo* experiments by Kierzkowski *et al*. [43] that identified a bias towards simpler leaf shapes in *Arabidopsis thaliana* and *Cardamine hirsuta*.

In particular, this study found that expression of the genes *SHOOTMERISTEMLESS* (*STM*) and *REDUCED COMPLEXITY* (*RCO*) together was necessary to generate the compound shape in *C. hirsuta* and sufficient to induce compound shape in *A. thaliana* (fig. 2a). Therefore, in this system, compound shape requires a more specific combination of gene expression than simpler shapes (e.g. lobed, unlobed).

**FIG. 2.**
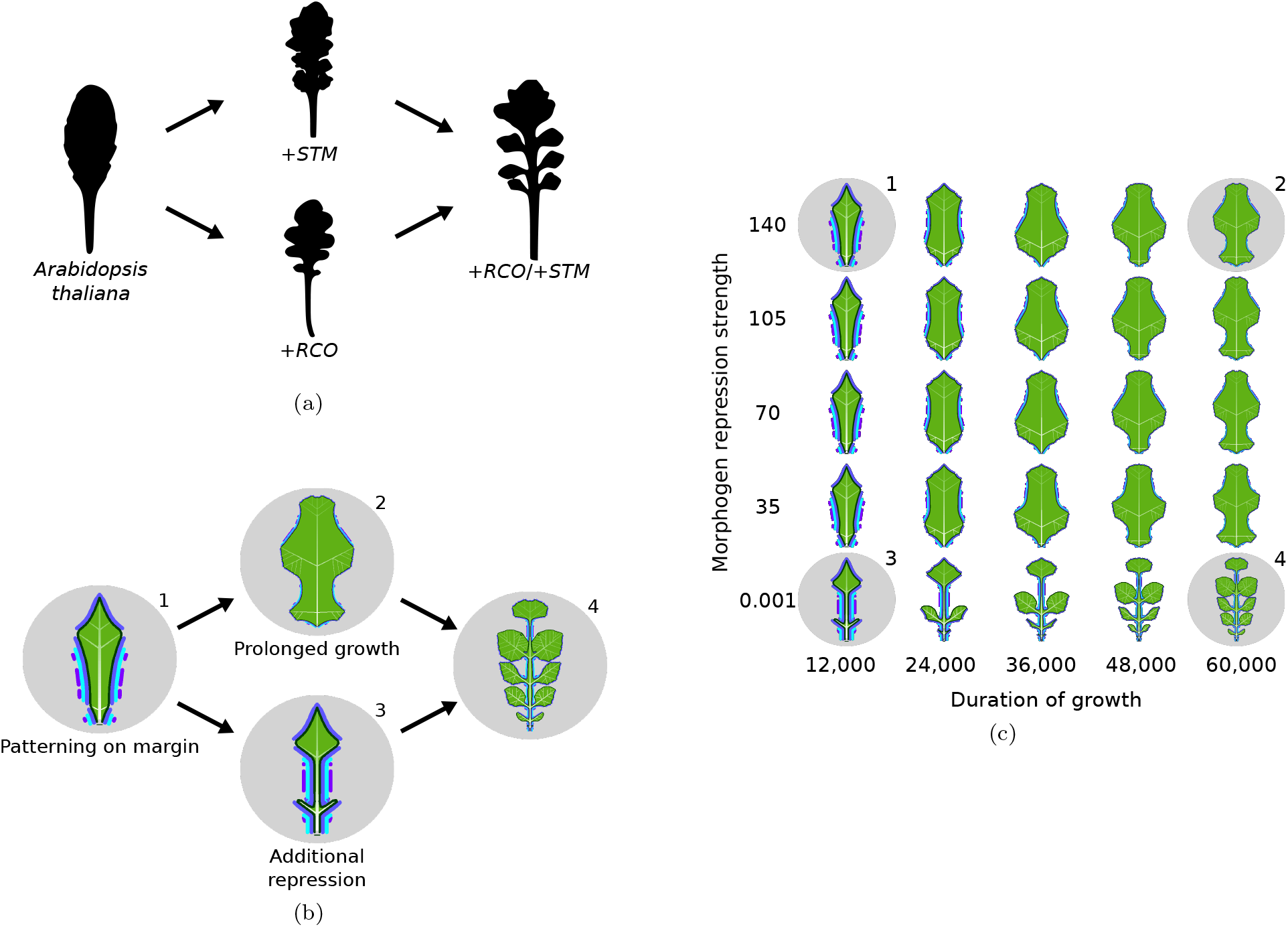
An experimentally motivated illustration showing that simple leaves are easier to generate than complex leaves. **(a)** Leaf silhouettes of wild-type *A. thaliana* and three mutants, which result in lobed (+*STM*, +*RCO*) and compound shape (+*STM* /+*RCO*) (reproduced from Kierzkowski *et al*. [43] with permission). Ectopic *STM* and *RCO* expression individually create a lobed shape, but together induce compound leaflets. **(b)** A reproduction of the Kierzkowski *et al*. [43] result using the Runions *et al*. [14] computational model of leaf development. +*STM* is approximated by increasing the duration of growth and +*RCO* by increasing the growth repression strength of one morphogen on the margin of the leaf. Morphogen presence is represented by the purple/blue lines outside the margin. **(c)** A morphospace showing the resulting shapes from different levels of the two parameters we use to approximate *STM* and *RCO*. The range shown for *morphogen repression strength* is the range in which biologically plausible shapes can be produced. The range for *duration of growth* is the range in which leaves can be generated in a reasonable timeframe. In the parameter files of the Runions *et al*. [14] model, the parameters we refer to as *morphogen repression strength* and *duration of growth* are given by FAIRING4 and FINALFRAME respectively.

While the Runions *et al*. [14] model was not designed to model this system specifically, we tested whether it could nonetheless reproduce the result from Kierzkowski *et al*. [43] by selecting two model parameters that control similar growth properties to the genes *STM* and *RCO*, total growth duration and morphogen repression strength respectively (fig. 2). In agreement with the experiments, we found that activating both parameters together was sufficient to produce a compound shape while activating each in isolation was not (figs. 2a and 2b).

Having reproduced the experimental phenotypes, we next used the model to explore the *STM* /*RCO* parameter space in more detail, systematically varying morphogen repression strength and duration of growth within their biologically plausible ranges (fig. 2c). We found that the majority of this space (20 of 25 parameter combinations) was occupied by simple, unlobed shapes, with only a small fraction representing more complex shapes (1/25 dissected, 4/25 compound—see Methods section IV D). This agrees with Kierzkowski *et al*. [43] who also found that a similar fine-tuning is required for compound leaves.

### C. Mutation in a computational model of leaf development is sufficient to reproduce phylogenetic bias towards simple leaves

The mechanism derived from the experiments on varying *STM* /*RCO*, shown in fig. 2, is one specific driver of developmental bias. To investigate a wider range of possibilities, we study the effect of random mutations for a broad set of Runions *et al*. [14] model parameters (table S1). We first generated a sample of initial leaves representing a diverse morphospace (figs. 1b and S6). We mutated the initial sample using a random walk algorithm with two different mutational schemes, MUT2 (alg. 1) and MUT5 (alg. S2). These algorithms work by iterating through model parameters one by one and perturbing the value by a small amount. We then automatically classified the resulting shapes into the same four shape categories (unlobed, lobed, dissected and compound) using a morphometric classifier (fig. 6). Of the 76,800 total leaves generated per simulation, we found that the proportions of unlobed, lobed, dissected and compound were 73.4%, 9.6%, 11.4% and 5.6% respectively for MUT2 and 76.1%, 7.2%, 11.4% and 5.4% respectively for MUT5. This shows that our results are robust to the two different mutational schemes. Both reflect the fact that unlobed leaves occupy a significantly greater proportion of model parameter space that produces viable leaves than more complex shapes do (see also Results section II D).

Regardless of the initial leaf shape, over the course of these random walks, the proportion of walks in an unlobed state increased to an equilibrium just below 80% while most other shape categories stabilise to a significantly lower proportion (figs. 3 and 8). These predictions lie within the margins of error of the phylogeny-derived CTMC model predictions (figs. 3 and 8), suggesting that mutation and development alone are, to first order, sufficient to predict the relative frequency of different leaf shapes over evolutionary time.

**FIG. 3.**
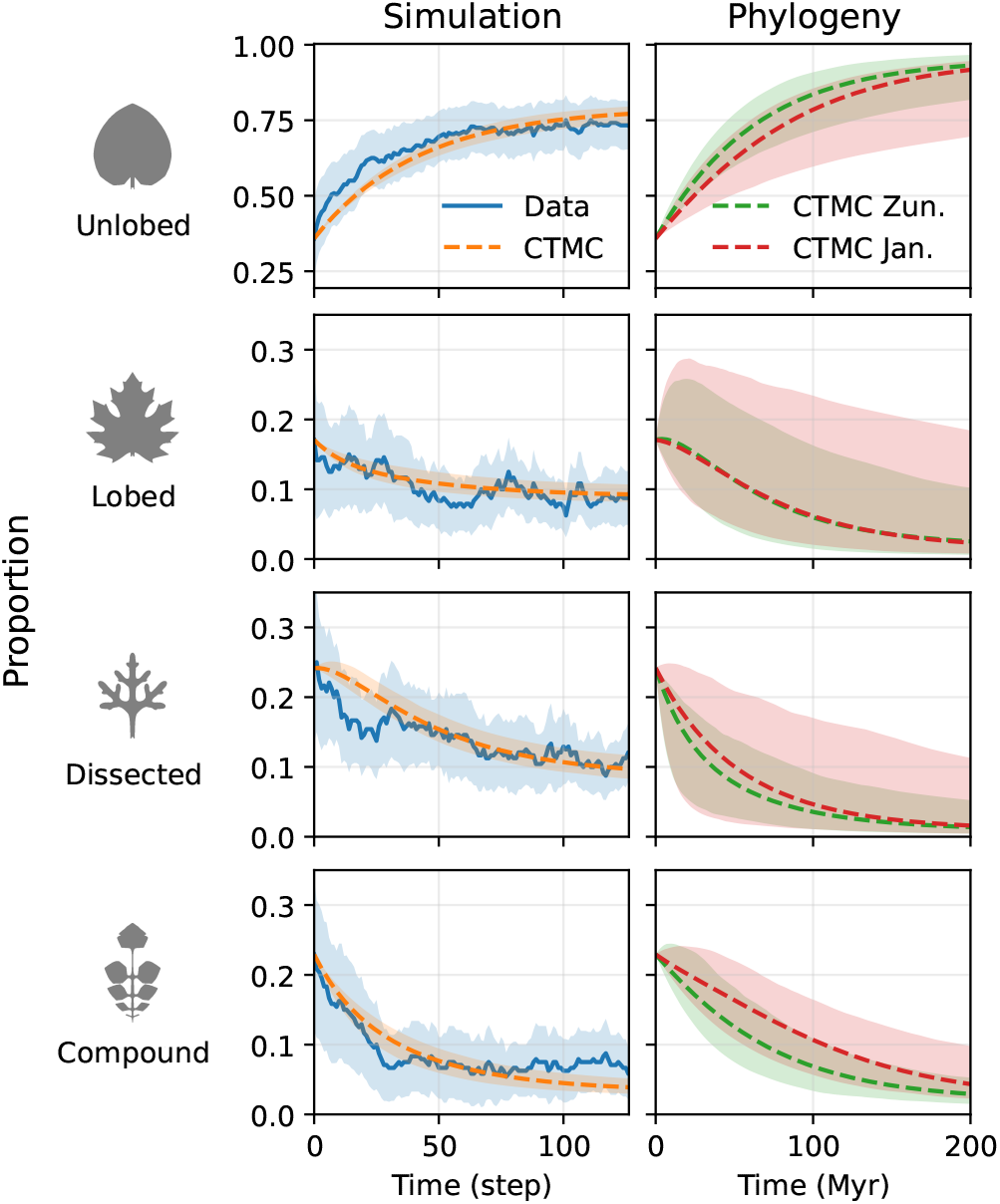
The proportions of each leaf shape over time as predicted by *in silico* mutagenesis and phylogeny. (Simulation) The solid blue line represents the mean proportion of random walk chains in each state at each time step (*±*1.96*×* standard error) in our simulation (MUT2 alg. 1). To these data, we fit a CTMC via maximum likelihood estimation (MLE) (dashed line) and MCMC with errors inferred from MCMC posteriors via Monte Carlo error propagation. Since step size is arbitrary, for comparison with the phylogenetic model, we adjust the x-limits for the simulation such that the CTMC predictions for both the simulation and phylogeny reach halfway from the initial distribution to the stationary distribution by approximately the same distance along the axis. **(Phylogeny)** We predict how the proportions of each leaf shape evolve from the same initial proportions used in the simulations, using the CTMC inferred from each phylogeny (green: Zuntini *et al*. [41] genus (fig. S9), red: Janssens *et al*. [40] genus (fig. S11)). Dashed lines represent the prediction using MLE transition rate values. Errors for all CTMC predictions were inferred from the transition rate MCMC posteriors using Monte Carlo error propagation (*N* = 500), and represent a 95% credible interval for each time step. We show these extrapolations for a time period equal to the approximate age of the angiosperm clade (200 Myr) [41].

To enable a more detailed comparison between simulation and phylogeny, we inferred the rates of shape transitions from our simulations. By estimating rates for all 12 possible (non-self) transitions, we gain deeper insight into the dynamics of shape change over time than by examining the proportions of the four shape categories alone. To do this, we fitted the same CTMC model used for the phylogenetic inference to the random walk data, to estimate the evolutionary rates between shape categories in the absence of processes other than mutation, using MCMC [44, 45]. In agreement with our phylogenetic inference, we found that transitions from unlobed occurred at a significantly lower rate than transitions to unlobed, giving a consistent net transition rate towards unlobed (figs. 1b and 7a). This robust agreement between relative rates derived independently from simulations and phylogenies suggests that the evolutionary preference for simple leaves observed across angiosperm phylogeny may primarily be driven by developmental bias instead of selection.

By contrast, for transition rates between complex leaf shapes (lobed, dissected and compound) (figs. 1 and 7a) there is less agreement both among the different phylogenies and between the phylogenies and the simulations.

These differences could be because selection has played a more prominent role in the transitions among more complex leaf shapes than it has for transitions from complex to simple. The model may also be less reliable at resolving relative rates of complex leaves, for which more detail needs to be specified. Fully resolving the discrepancies for transitions between more complex leaf shapes would require reducing the error associated with the phylogenetic rate estimates, which in turn would necessitate significantly more leaf shape data.

### D. Complex leaves occupy a smaller volume in parameter space and are closer to simple leaves, but not vice versa

To further investigate the mechanisms behind the bias towards simple leaves, we explored—much as we did for the experimentally inspired study in fig. 2c—the relative volumes in parameter space that different shapes occupy. Projecting the parameter values of all random walk outputs into a 2D space via principal component analysis (PCA) reveals several possible factors affecting relative rates (fig. 4). First, it shows that the four shape categories occupy areas of different sizes (fig. 4a). The simplest category, unlobed, occupies by far the greatest area (fig. 4b), suggesting that the bias towards simple forms in our simulations can be partially explained because, as argued before (fig. 2), more parameter combinations map to simple leaves than complex leaves. We find that this pattern also holds for PC3-PC4 (fig. S22e) and PC5-PC6 (fig. S22f).

**FIG. 4.**
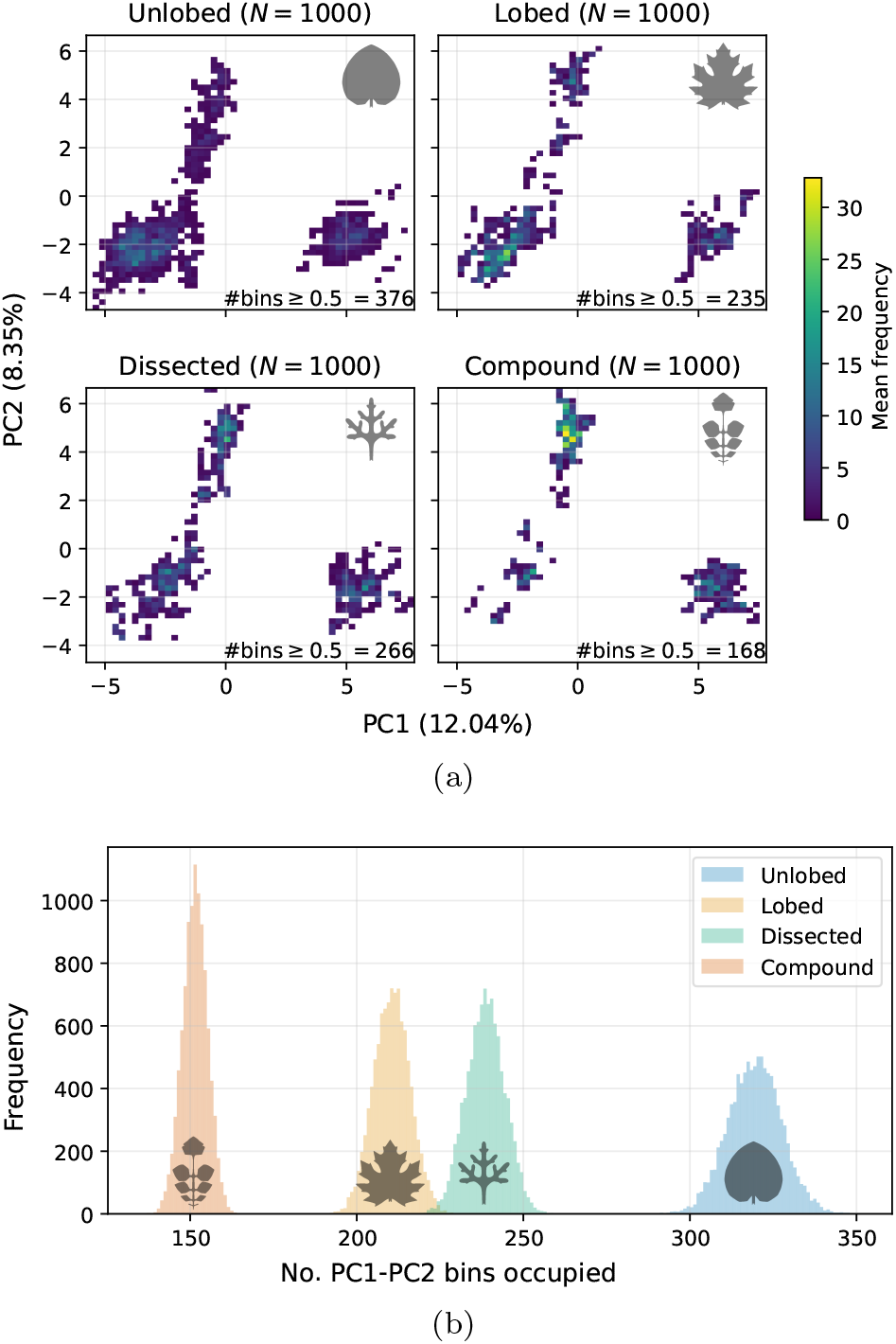
Simpler leaves occupy a greater volume than complex leaves in model parameter space. **(a)** A 2D histogram of model parameter space given by the 2 principal axes from a principal components analysis (PCA) of a sample of 4000 leaves from each shape category from one of the simulations (MUT2). Mean bin occupancy is calculated via a bootstrapping approach, by sub-sampling with replacement *N* = 1000 leaves from each shape category for 10000 iterations. Bins are shaded if the mean occupancy across these iterations is *≥* 0.5. This illustrates how unlobed leaves occupy a larger region of model parameter space than more complex shapes and that this larger space also contains the majority of more complex leaves. Moreover, more complex leaves are concentrated within a much smaller area than unlobed leaves. **(b)** The no. PC1-PC2 2D histogram bins occupied by each leaf shape in 10000 bootstrap samples (*N* = 1000) of a balanced sample of MUT2 (4000 per shape category). Bins with at least 1 leaf present are counted. This reveals that the unlobed category occupies by far the largest area in this 2D projection of parameter space. We find the same is true for other PCs (fig. S22).

Moreover, the vast majority of lobed, dissected and compound leaves are contained within a broader area occupied by unlobed in the 2D parameter space (fig. 4a), suggesting that more complex leaves are close to simpler leaves in parameter space but not always vice versa. This provides another mechanism making transitions from complex leaves to simple leaves easier than the reverse. We also observe this same pattern for PC3-PC4 (fig. S22b) and PC5-PC6 (fig. S22c).

Finally, the parameter space of unlobed leaves appears to be much more connected than the space of more complex leaves. Constructing a nearest neighbour graph (no. neighbours = 6) from the PC1-PC2 parameter space reveals that unlobed separates into far fewer connected components (3) than lobed (131), dissected (96) and compound (235) (fig. 9). Repeating this analysis in the full-dimensional parameter space also yielded fewer connected components for unlobed than for the other categories. This may further contribute to the bias in transition rates towards unlobed, as less connected spaces would be more frequently exited by random walks than more connected spaces.

Our results show that the finding from experiments [43] and manual parameter searches (fig. 2c)—that simpler leaves occupy a larger volume within the parameter space of the developmental programme—extends to the Runions *et al*. [14] model more generally. This volume disparity, in addition to other properties including the degree of overlap between shape categories and the connectedness, may account for much of the bias towards unlobed we see in our simulations.

## III. DISCUSSION

We set out to test the hypothesis that mutation alone was sufficient to produce the higher transition rates towards simple leaves compared to complex leaves observed in angiosperm phylogeny [39]. By performing *in silico* mutagenesis, we found that non-adaptive processes alone produce greater transition rates to simple unlobed leaves than to complex leaf shapes such as lobed or compound (figs. 1b and 3). We also reproduced the observation from angiosperm phylogeny that unlobed leaves have evolved at a faster rate from complex leaves than vice versa [39], using larger and more recent trees [40, 41] and new classification data (figs. 1a and 3). Together, this suggests that a developmental bias towards simple, unlobed leaves can explain this macroevolutionary pattern in flowering plants.

Our finding is consistent with the growing body of evidence that developmental bias can play a significant role in evolutionary outcomes for morphological traits [4–11] The particular bias towards simplicity is consistent with recent work arguing for a generic bias towards descriptional simplicity in genotype-phenotype maps [8]. Johnston *et al*. [8] provided examples of this bias strongly affecting RNA secondary structure, where simpler structures are much more likely to be observed in nature than complex ones (see also [46]) and for self-assembling protein quaternary structure, where this bias helps explain the prevalence of highly symmetric forms in the Protein Data Bank (PDB). Furthermore, this bias has been predicted to hold more widely for input-output maps [47]. This generic argument implies that the simplicity bias observed in molecular phenotypes may extend to the level of macroscopic phenotypes, which we show here for leaf shape, but also potentially to a much wider set of morphological phenotypes. More specifically, our results suggest that this bias towards simplicity may hold generically when development uses conserved mechanisms and GRNs where complex phenotypes need tighter parameter specification than simpler phenotypes.

Our study also opens up a number of future directions for studying the evolution of leaf shape. First, it would be valuable to repeat similar simulations in alternative models of leaf development [43, 48, 49]. The Runions *et al*. [14] model is uniquely suited to macroevolutionary studies such as this one, as it captures key conserved elements of angiosperm leaf development while generating a large morphospace, but, if the same bias towards simplicity was observed in other models, this would provide further evidence that it is a general property of leaf development. Second, it would be illuminating to conduct similar *in silico* mutagenesis, but classify the resulting leaves using a continuous classification system instead of discrete categories, such as directly measuring margin complexity using a compression algorithm on the leaf silhouette [50]. This could help identify more subtle biases between different leaf forms, including teeth/serrations, which we do not analyse here, but which the model can produce (figs. 1b and S6). Third, when inferring transition rates from phylogeny, we were limited by the availability of herbarium data, the resolution of phylogenetic trees and the rate at which herbarium leaves could be manually classified. To more accurately resolve transition rates between more complex shape categories, it would be useful to repeat this study with larger phylogenies. This would likely require an automated method for herbarium shape classification. Finally, the methodology presented in this study—combining phylogenetic analysis with evolutionary and developmental computational models—could be used to study the evolution of other traits where a reliable developmental model exists, and phylogenetic data are available (e.g. [5]). This would greatly expand our understanding of how well development can predict trait evolution across phylogeny.

## IV. METHODS

### A. Simulating leaf development

We use a well-established model by Runions *et al*. [14] to approximate the developmental programme underpinning angiosperm leaf shapes. In this model, leaves are represented in 2D as three components (figs. 5a and 5b). First, an open tree structure with growing edges to which new edges are added, representing the vasculature; second, a polygon with morphogen presence or absence at each vertex that deforms according to the growing vasculature and instructs the initiation of new veins, representing the margin; third, a passive triangular mesh representing the lamina that fills the space inside the margin.

**FIG. 5.**
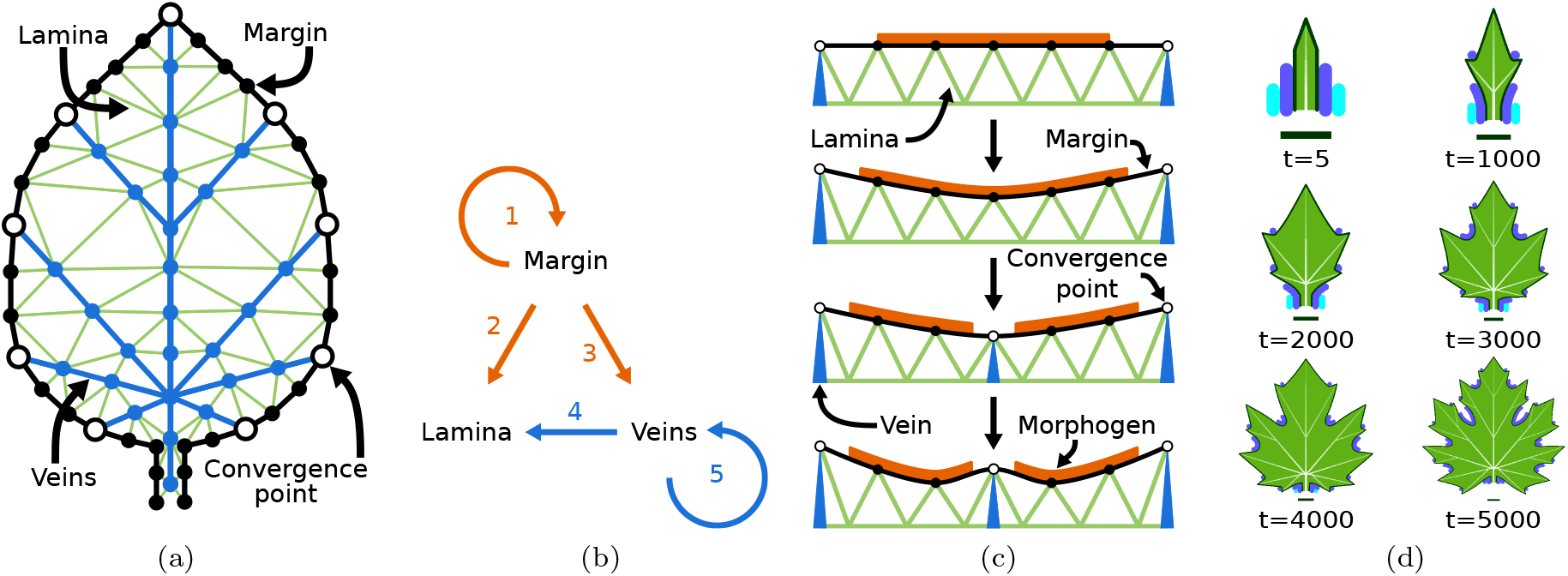
A description of the leaf model from Runions *et al*. [14], highlighting the three key components of leaf development and their interactions. **(a)** Model components illustrated in leaf form (inspired by Runions *et al*. [14] fig. 3). Margin in black, veins in blue, lamina in green, convergence points in white with black outlines. **(b)** The three components of the model with arrows indicating interactions that affect growth. (1) Existing morphogens and convergence points influence where new ones will form. (2) Convergence points and morphogens influence local growth rate. (3) Convergence points induce new veins. (4) Veins determine local growth direction. (5) Existing veins influence the growth of new veins. **(c)** Zoomed view of the margin, showing how growth of the vasculature and patterning on the margin interact to produce protrusions and sinuses. Veins (in blue) extend, pushing the margin out; however, a morphogen (in orange) inhibits webbing between the veins, causing a sinus to form. This increases the distance between the two convergence points, leading to the insertion of another convergence point between the two that excludes the morphogen from its vicinity. Existing vasculature connects to the convergence point and specifies a new axis of growth. **(d)** Different growth stages of a model leaf. The dark blue morphogen inhibits webbing and convergence point formation; the light blue morphogen delineates the petiole. The value of *t* denotes the iteration of the simulation.

Many similar models capture a narrow range of shape diversity—for example, variation within *A. thaliana* [48, 49]. On the other hand, the Runions *et al*. [14] model captures a broad spectrum of leaf shapes within a single parameter space. This enables the inference of transition rates between highly divergent shapes such as unlobed and compound. However, it also introduces challenges as the genetic basis of leaf shape differs among lineages and remains poorly resolved for many non-model species. To address this, the Runions *et al*. [14] model is formulated at the level of developmental mechanism rather than specific genes. It is therefore important to understand that this model is limited in what it can tell us about leaf development and evolution at the level of specific genes. Growth in the Runions *et al*. [14] framework is a product of two key developmental mechanisms. The first is morphogen patterning on the leaf margin. The margin—the tissue at the adaxial-abaxial boundary—has long been recognised as a critical site for patterning during leaf development. For example, auxin patterns marginal outgrowths in *Solanum* [51], *A. thaliana* [48] and *C. hirsuta* [52]. The products of genes such as *CUC2* in *A. thaliana* [48] or *RCO* in *C. hirsuta* [43] also localise to the margin and determine whether these auxin-driven outgrowths form serrations, lobes or leaflets. The closely related *CUC3* and *NO APICAL MERISTEM* (*NAM*) genes localise to leaflet primordia boundaries, and their down-regulation reduces serration and lobing in diverse lineages, namely *Aquilegia caerulea, Pisum sativum, Cardamine hirsuta, Solanum lycopersicum* and *Solanum tuberosum* [53]. Although the role of auxin in patterning protrusions is highly conserved, many other developmental regulators are not conserved across phylogeny. For example, *RCO* is only present in the Brassicaceae family [54] and therefore plays no role in determining leaf shape in other compound lineages such as *Solanum*. Furthermore, there are likely many other genes regulating leaf shape in non-model species that are currently unknown. The model accounts for this variation and uncertainty by not assuming a conserved identity for the morphogens. Both the number of morphogens present and their effect on growth can vary according to their associated parameter values. What is assumed to be conserved is the broader principle that growth-modifying morphogens form patterns along the margin.

The second key mechanism that the model incorporates is the alignment of local growth direction with veins. This is consistent with the telome theory [55], according to which leaves first evolved by the webbing of tissue between shoots, as well as a number of lines of empirical evidence. At the tissue level, deeply dissected leaves (e.g. *Pelargonium carnosum*) necessarily align growth directions with main veins and variegated leaves also show growth orientations that track major veins [14]. At the molecular level, auxin has been shown to pattern veins [56, 57] as well as marginal outgrowths [48, 51, 52] in model species, consistent with the model assumption that vein growth drives margin growth. In the Runions *et al*. [14] model, consistent with observed patterns of auxin efflux carriers in *A. thaliana* [58], new veins are inserted between the current vasculature and special points on the margin known as “convergence points”. New convergence points form between old convergence points depending on the distance between old points and the effect of any morphogens present (fig. 5c). This is an extension of Hofmeister’s rule [59] for leaf primordium emergence in the shoot apical meristem, where new primordia emerge in the largest space between existing primordia. Some properties of veins are not captured by the model, specifically closed reticulate and parallel vein architectures. This is partly because there is some uncertainty surrounding the mechanisms by which these architectures form [57, 60], but we acknowledge this as a limitation of the model.

More broadly, much of our understanding of leaf development derives from a small number of model species. These have revealed some conserved genes and mechanisms [61], but many details are lineage-specific [54]. Consequently, some unavoidable uncertainty remains in how well the Runions *et al*. [14] model captures global trends.

### B. Generating the initial sample

Since a range of different forms are contained within each shape category, we began our simulations from leaves drawn from an initial sample representing a wide variety of morphologies (16 unlobed, 8 lobed, 11 dissected, 13 compound) (figs. 1b and S6). This sample was generated using a variety of methods. Some leaves were taken from Runions *et al*. [14]; others were generated by tuning parameters by hand and some were taken from the output of random walks on leaves from Runions *et al*. [14].

### C. *In silico* mutagenesis

To simulate mutation in our sample, we used a constrained random walk algorithm. The algorithm samples the parameter space of biologically plausible leaf shapes surrounding the starting leaf by making a small change to one of the parameters, checking the validity of the resultant leaf (table I), and then moving on to another parameter (see fig. S8 for examples of rejected leaves). We chose this approach over other methods, such as uniform sampling or Latin hypercube sampling, as the biologically relevant ranges of each parameter were unknown. Random walks have been shown to cause some biases towards high-robustness regions of parameter space in discrete parameter spaces represented as networks [62], which in this case would result in the underrepresentation of invalid leaves. We have no reason to assume that this would systematically bias the rates between valid leaves, but this cannot be fully excluded. For each of the 48 leaves in our initial sample (fig. S6), we ran 5 random walks for 320 steps, generating a total of 5 × 48 × 320 = 76,800 per simulation. We allow 100/122 model parameters to vary throughout the simulations (table S1). Not all parameters are included, as some are not relevant to leaf shape (e.g. they control how the image is rendered).

**TABLE I.**
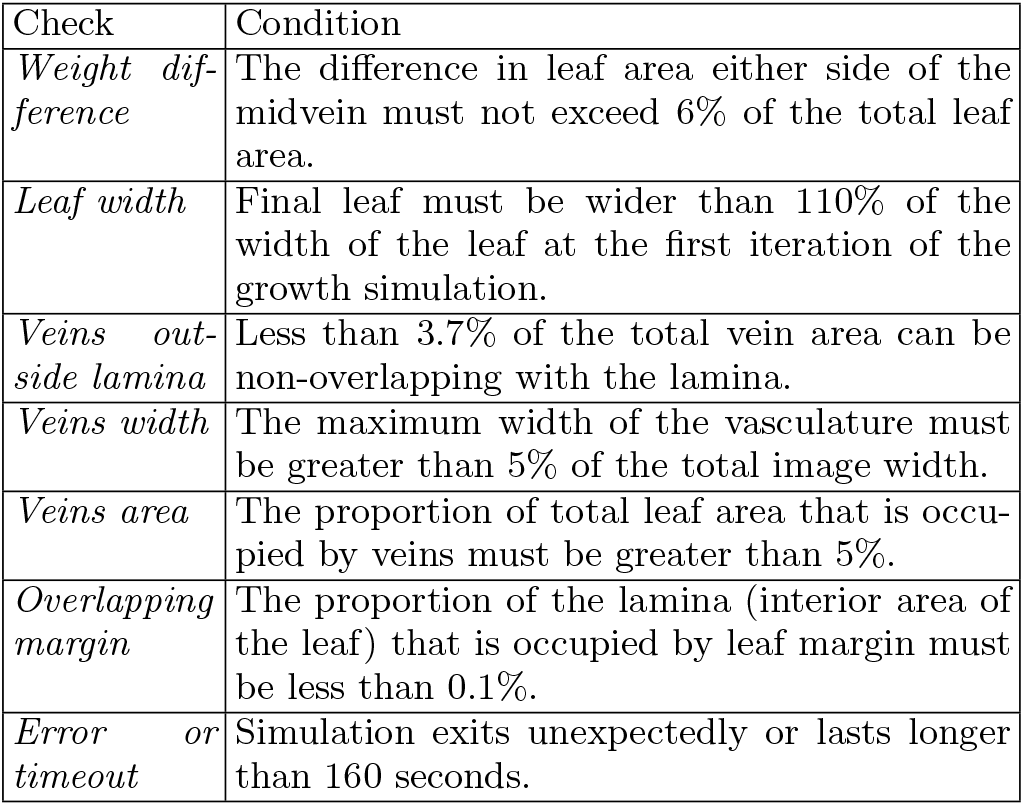
Random walk constraints. The only type of selection we apply during the walk is stabilising selection for physically plausible leaves. See fig. S8 for examples of rejected leaves. *Weight difference* (fig. S8a) ensures that the midvein and petiole are roughly in line with the centre of mass of the leaf. *Leaf width* (fig. S8b) ensures that leaves which stop growing after the first few stages of the model simulation are rejected. *Veins outside lamina* (fig. S8c) prevents leaves forming whose veins grow outside the lamina. *Veins width* (fig. S8d) and *Veins area* (fig. S8e) ensure that leaves that generate without adequate vasculature are rejected. *Overlapping margin* (fig. S8f) rejects leaves with ingrown margins. *Error or timeout* – some parameter combinations result in the simulation exiting unexpectedly or taking a very long time to complete, posing an additional constraint.

| Check | Condition |
| --- | --- |
| *Weight difference* | The difference in leaf area either side of the midvein must not exceed 6% of the total leaf area. |
| *Leaf width* | Final leaf must be wider than 110% of the width of the leaf at the first iteration of the growth simulation. |
| *Veins outside lamina* | Less than 3.7% of the total vein area can be non-overlapping with the lamina. |
| *Veins width* | The maximum width of the vasculature must be greater than 5% of the total image width. |
| *Veins area* | The proportion of total leaf area that is occupied by veins must be greater than 5%. |
| *Overlapping margin* | The proportion of the lamina (interior area of the leaf) that is occupied by leaf margin must be less than 0.1%. |
| *Error or timeout* | Simulation exits unexpectedly or lasts longer than 160 seconds. |

To test the robustness of our results, we repeated the simulation with 2 different mutational schemes. MUT2 (alg. 1) iterates through the parameters in a random order, and attempts to change the parameter by a value selected at random from an array of numbers randomly generated at 3 different orders of magnitude. MUT5 (alg. S2) is the same as MUT2 except that each mutation is additionally scaled by 10% of the current parameter’s range within the initial leaves (fig. S6). The aim here was to provide some way of accounting for the biologically relevant sampling range.

### D. Shape classification

To sort the leaves into each of the four shape categories (unlobed, lobed, dissected, compound), we developed a computer-vision program (fig. 6) to reduce error from subjectivity. It finds the coordinates of key landmarks in the leaf image, extracts morphometric measurements and assigns a category based on threshold values of these measurements (eq. (1)). These morphometric measurements were also used to check if leaves were valid (table I) in the random walk algorithm.

#### Algorithm 1

Random Walk MUT2

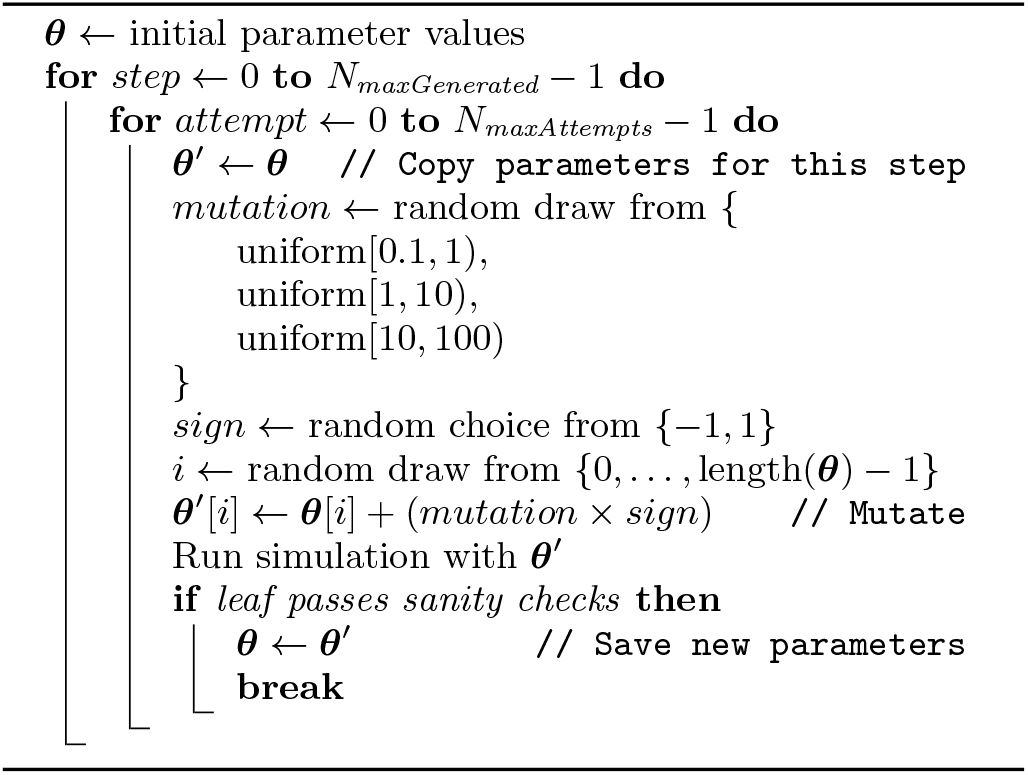

First, the main visual features are extracted from the leaf image, including silhouette, margin, lamina and veins. The coordinates of each pixel along the edge of the silhouette (the contour) are extracted. A reference point is defined in the centre of the midvein at the point where it first branches outwards into lateral veins (fig. 6b). Then, the distance between each point on the contour and this reference point is calculated (fig. 6c). The local maxima and minima in this distance array that pass a certain prominence threshold are determined using `find_peaks` from the scipy.signal package with `distance=10` and `prominence=25`. These pertinent extrema represent estimates for the coordinates of the tips and sinuses of any protrusions in the leaf margin (fig. 6b shown in red and blue). Because of the prominence threshold, small protrusions such as serrations are excluded.

**FIG. 6.**
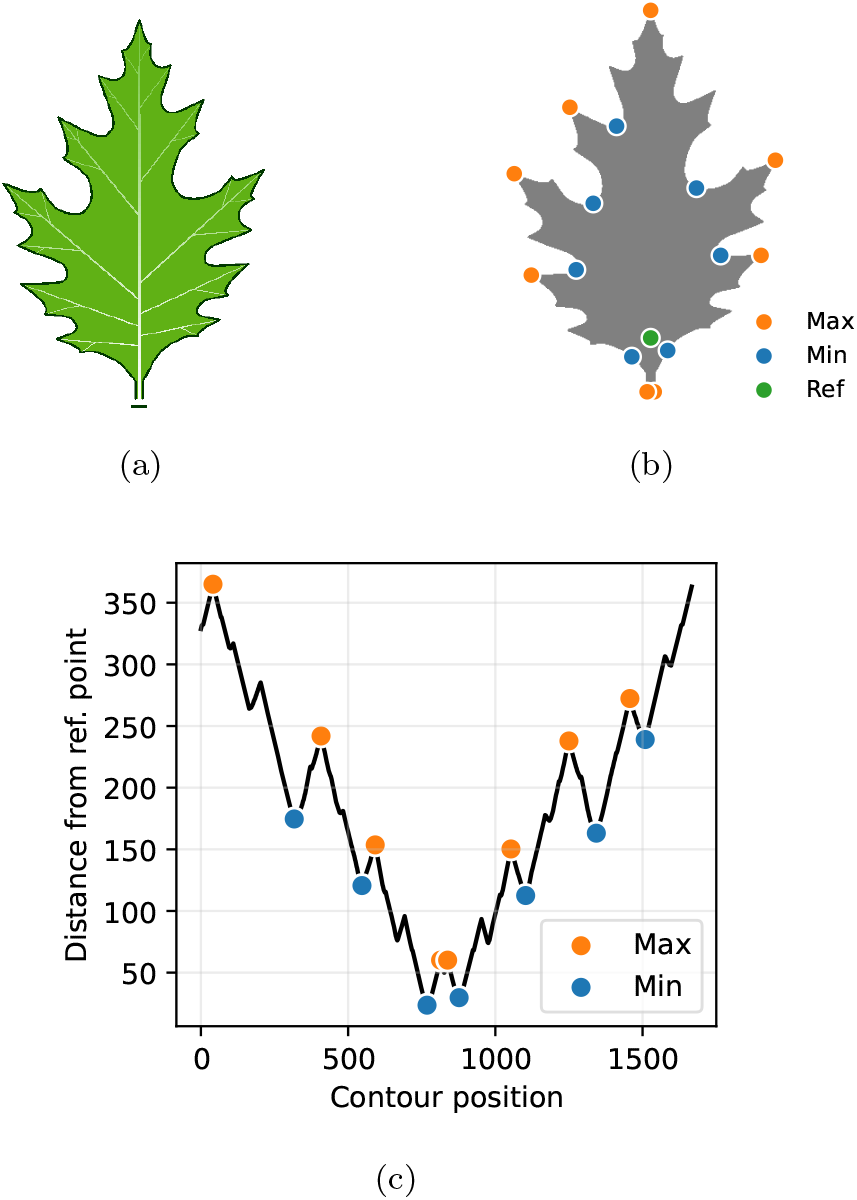
A summary of the morphometric classifier. **(a)** Example lobed leaf. **(b)** Silhouette with local maxima in orange, minima in blue and reference point in green. **(c)** Graph of distances of each contour point to the reference point. Local maxima shown in orange, minima in blue.

From the coordinates of the local minima, maxima and the reference point, we calculate a number of morphometric measurements which are then used to classify the leaf. Three measurements are used: *ϵ*, the total no. pertinent local maxima and minima; *α*, the average angle formed by each local maximum and its two adjacent minima; and *ω*, the average horizontal distance between each pair of local minima, closest to the midvein, that share the same vertical coordinate (measured in pixels). The definitions for each of the shape categories are as follows:

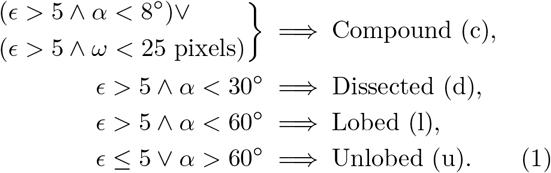

Any leaf that fails to satisfy any of these conditions is labelled as “unclassified”. However, there were no unclassified leaves in either the MUT2 or MUT5 simulation.

Leaf shape is a continuous trait. Thus, the boundaries between discrete categories, such as those in eq. (1), are arbitrary. This makes it difficult to assess the accuracy of the classifier algorithm. Nevertheless, we wanted to have some measure of how closely it matches classification by eye. Therefore, we manually classified 200 leaves randomly selected from one of the simulations, with an approximately equal number from the output of each initial leaf. 78.5% of the manually ascribed labels matched those assigned by the classifier (see fig. S7 for further details). For comparison, we fit a machine learning algorithm (Histogram-based Gradient Boosting Classification Tree from the scikit-learn Python package) to a sample of 500 manually labelled leaves, sampled in the same way as the set of 200. The algorithm was trained on the distance of 200 equally spaced points along the leaf contour to the reference point for each leaf (fig. 6c). When used to predict shape for the same sample of 200 leaves used for the threshold classifier, it returned an accuracy of 73.0%. We then repeated the machine learning approach using morphometric data rather than contour data, which returned an accuracy of 79.5%. This suggests that the threshold classifier is a good enough match with subjective shape classification compared to other classification methods. It also demonstrates that the morphometrics used to construct the original classifier are useful features when classifying shape as unlobed, lobed, dissected or compound. The two machine-learning methods agreed on 77.0% of the labels.

### E. Inferring evolutionary rates from random walk data

To test the model for bias in leaf shape transition rates upon mutation, we ran 5 random walks on each of the 48 initial leaves, for 320 steps, generating a total of 76,800 leaves for each of MUT2 and MUT5. Using the classifier (fig. 6), we assigned each leaf the category unlobed, lobed, dissected or compound (eq. (1)). Evolutionary rates between shape categories were inferred by fitting a continuous-time Markov chain (CTMC) to the labelled walks.

This model is defined by a matrix of transition rates ***Q*** between each state—unlobed (*u*), lobed (*l*), dissected

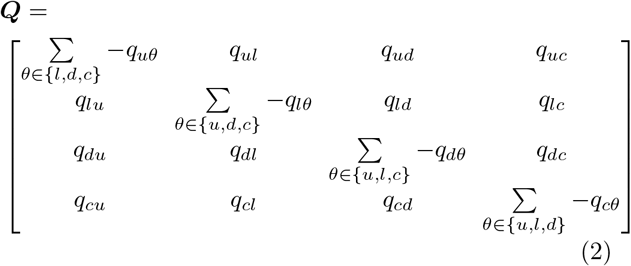

where the diagonal elements must be equal to the negative sum of all other elements in the same row [63].

The model assumes that at any given instant, over a continuous-time interval, the system is in one of these four states. The probability *p*(*i, j, t*) of transitioning between state *i* and *j* depends only on the current state, the transition rate matrix ***Q*** and the time *t* until the end of the transition. The matrix of all possible transition probabilities ***P*** (*t*) relates to ***Q*** and *t* as follows [44, 63]

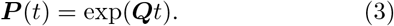

Therefore, the probability of a transition from state *i* to *j* over time *t* is

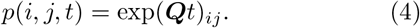

The probability of a set of observed walks *x* = {*x*_0_, *x*_1_, …, *x*_*W*_ }, where each walk contains a sequence

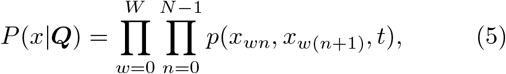

where *w* and *n* are walk and step indices, respectively. We assume *t* = 1 for every step, as over the course of the entire random walk, the average step size in parameter space will be approximately constant.

Rates were inferred from the random walk simulation data by finding the maximum of this function, using MCMC with the emcee package for Python [64]. We ran 24 chains for 5,000 steps with uniform priors between 0 and 0.1 for all transition rates. Chains were thinned by removing all but every 10th step (figs. S15 and S16) and the first 2,500 were discarded to obtain the posterior distribution. We took the mean of the posterior for each transition as the estimate for each transition rate in fig. 1. As an additional check, we also inferred the rates via maximum likelihood estimation (MLE) and found that they converged with the MCMC estimates (figs. 7a and 7b).

**FIG. 7.**
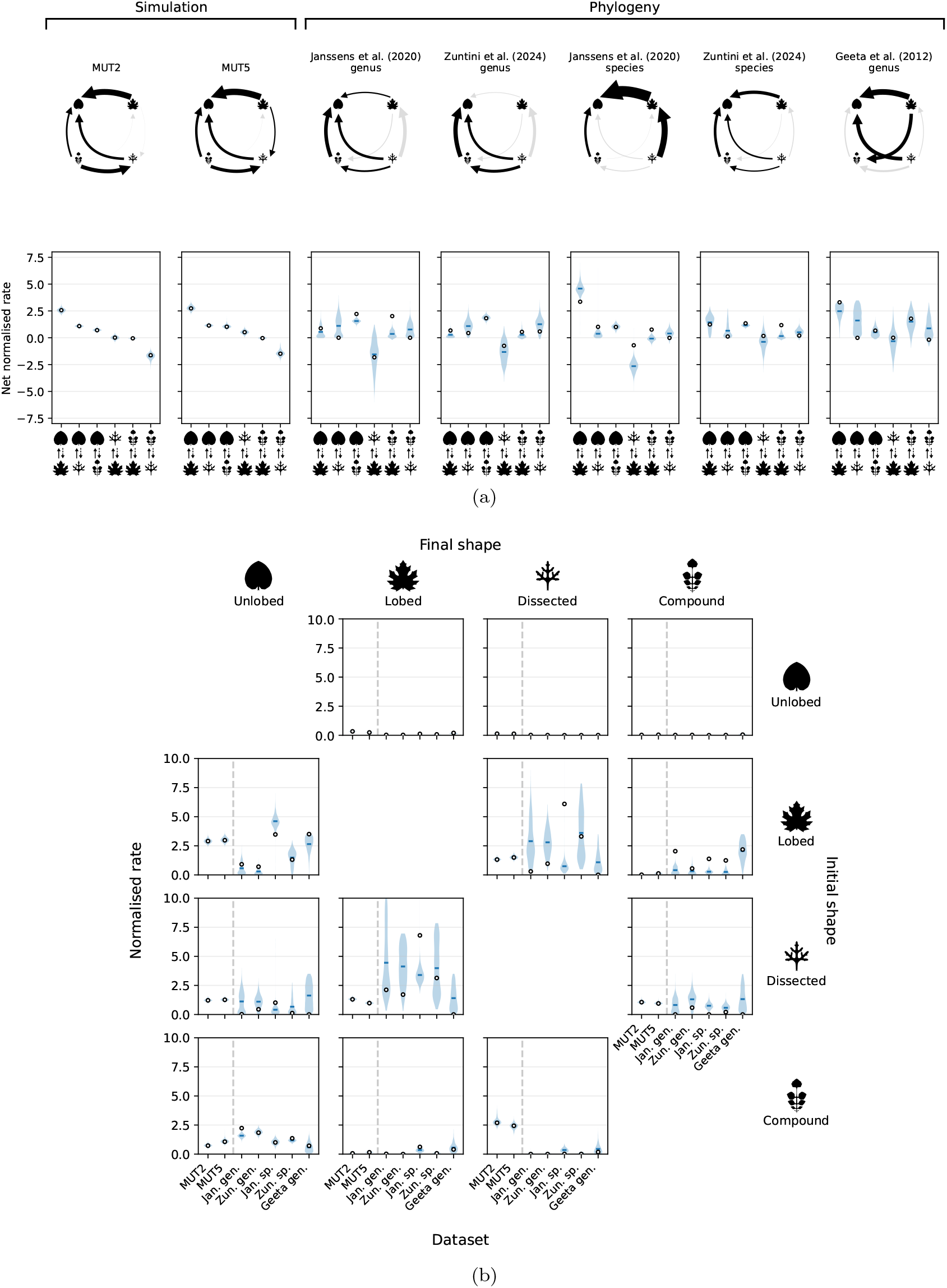
Further details of transition rates from phylogenies and simulations. **(a)** Net normalised transition rates between the 4 leaf shape categories, inferred from both simulation and phylogeny. (Top row) Arrow thickness and direction represent the mean net normalised transition rate between each pair of leaf shape categories. Black arrows represent transitions where 0 does not fall within an equaltailed 95% credible interval for the net normalised posterior, and grey where it does. (Bottom row) Posterior distributions for the net normalised rate of all transitions. Solid bars indicate the mean. The net rate is calculated by subtracting the transition given by the dashed arrow from the transition given by the solid arrow in the x-axis labels. **(b)** Normalised transition rates between different leaf shapes, inferred from both *in silico* mutagenesis (left of dashed line) and phylogeny (right of dashed line). Each subplot represents a different transition, from the shape shown on the vertical figure axis to the shape shown on the horizontal figure axis. These rates were normalised by dividing by the mean transition rate estimate within each dataset (eq. (6)).

**FIG. 8.**
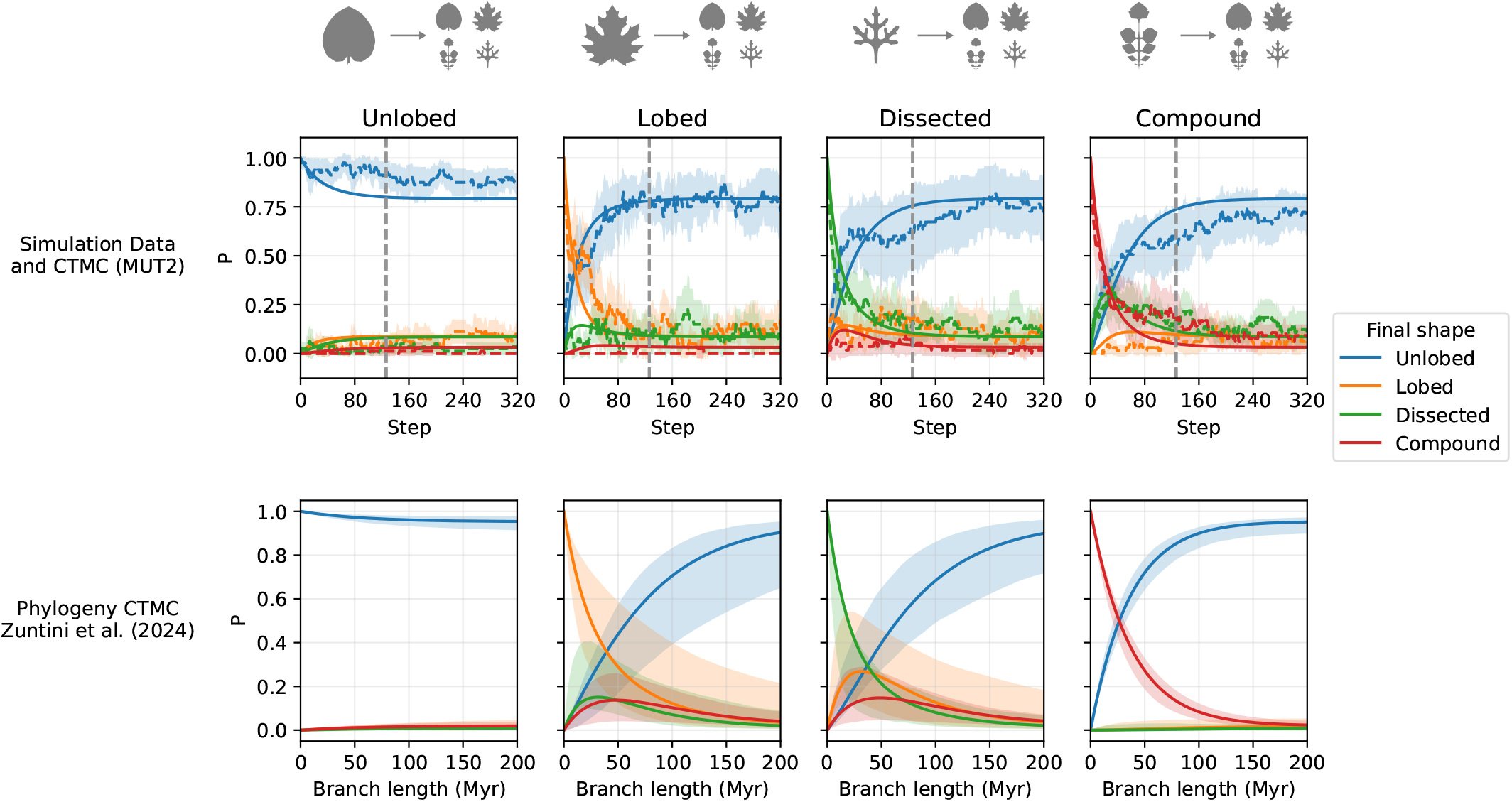
Comparing *in silico* mutagenesis and phylogeny for the probability of each shape transition over time, separated out by starting shape. The column header gives the initial shape and the line colour gives the final shape. **(Top row)** The dashed line represents the proportion of walks from each unique initial leaf (fig. S6), averaged over all initial leaves in each state. The error bands represent *±*1.96*×*standard error for this average. The solid lines represent the CTMC model fit (MLE) to the output of the random walks. The vertical, grey, dashed line represents the estimated simulation step that is equivalent to *t* = 200 Myr for the phylogeny CTMC. This is calculated as described in fig. 3, and is also the simulation x-limit in fig. 3. We show all 320 simulation steps here for a more detailed summary of the simulation trajectories. **(Bottom row)** The predicted transition probabilities given by the CTMC model fit to the labelled Zuntini *et al*. [41] genus tree (fig. S9). The solid line represents the MLE fit and the errors are inferred from MCMC posteriors via Monte Carlo error propagation (*N* = 500) as per fig. 3. Note the good qualitative agreement between the phylogenetic and simulation dynamics, especially for the (blue) data related to transitions to/from unlobed leaves.

**FIG. 9.**
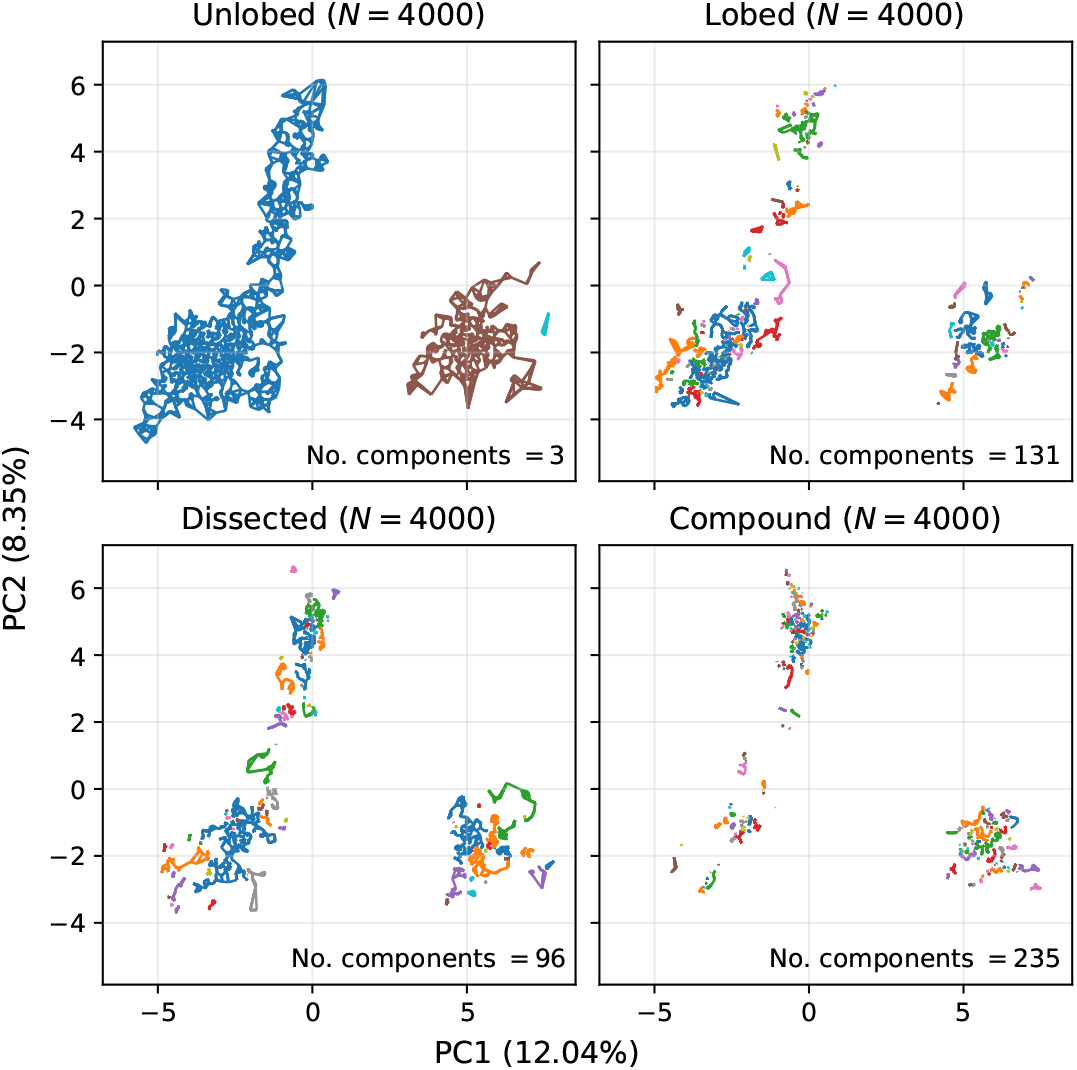
The nearest neighbour graph of parameter PCA space for each shape category highlights how the unlobed space is the most connected, followed by dissected, lobed and then compound. Complex leaves subdivide into many more connected components than unlobed. This pattern is also observed when the nearest-neighbour graph is constructed in the full-dimensional parameter space, where the numbers of connected components are 146, 273, 255 and 315 for unlobed, lobed, dissected and compound leaves, respectively. Thus, the greater connectedness of the unlobed parameter space is not an artefact of the PCA projection. The number of nearest neighbours used is 6. Colours highlight different connected components; however, since the number of unique colours is only 10, this under-represents the true number of connected components for lobed, dissected and compound. Thus, the colours here are intended only as a visual guide.

Because the inferred rates depend on the unit of time, we needed to normalise the transition rates to compare with the rates inferred from phylogeny. We did this for each mutational scheme by dividing by the mean of all transition rate estimates *q*_*ij*_. We define a normalised rate as

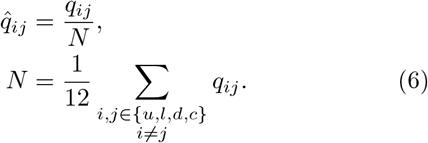

### F. Inferring evolutionary rates from phylogeny

Although Geeta *et al*. [39] had already inferred rates between these 4 categories with a phylogeny of 561 taxa, we wanted to test how robust these results are. Those results used a phylogeny that is now 25 years old [65], and there are now much larger (e.g. Zuntini *et al*. [41]—9,506 species) and more reliable phylogenies available. To do this analysis, we combined 2 phylogenetic trees [40, 41] with leaf images from the Naturalis herbarium [34]. We first filtered all digitised records to just those from angiosperm families as per APG IV [66]. We then removed all but the first record for each species.

To see how the rates are affected by looking at the species-level versus the genus-level, we generated 2 trees for each original tree, one with species at the tips and one with all but one tip per genus removed. For the species-level trees, we took the intersection between the filtered herbarium data and the full trees. If more than 3,100 tips remained, we sampled at random 3,100 tips, to allow for manual classification within reasonable time limits. This gave a combined set of 5,552 leaf images to classify for both species trees. Notably, the intersection between the herbarium data and the Zuntini *et al*. [41] species tree gave 3,057 taxa. Since this was less than 3,100, no further sub-sampling was required, and so unlike all the other trees in this analysis, the Zuntini *et al*. [41] species tree represents the raw intersection between the original tree and the herbarium. For this phylogeny, our genus tree and our species tree should both have similar levels of reliability. For the Janssens *et al*. [40] phylogeny, we trust the genus-level tree more than the species-level tree, because the overlap with the herbarium data was not as good at the species-level. For example, taxa were more heavily concentrated in a smaller number of angiosperm families, compared to the other trees (fig. S14). Moreover, we sub-sampled down to 3,100 taxa from 16,436 in the raw intersection to make manual classification feasible—something not required for the other trees.

For the genus-level trees, we further sub-sampled the angiosperm filtered herbarium data to 200 records per family, or the maximum if fewer than 200 records were present. The purpose of this step was to sample more evenly over phylogeny, accounting for potential biases in the herbarium. We then took the intersection between genera in this sample and each of our phylogenies and randomly selected one species from within each genus that was present in the herbarium to use as the label.

This gave a combined set of 3,359 leaf images to classify across both genus trees.

These images were then used to classify each species as one of the 4 categories (unlobed, lobed, dissected or compound) or ambiguous. This was done by eye, relying on a sample of model leaves and their morphometric classifier output for reference (figs. 1b and S6). After classification, all species labelled as ambiguous were removed, leaving an unambiguously labelled phylogeny (table II and figs. S9 to S13).

**TABLE II.** The number of taxa in each shape category for each labelled tree used in our inference. Columns with multiple shape categories (e.g. ld) represent taxa which could fit into any of the listed categories.

| Tree | Unlobed (u) | Lobed (l) | Dissected (d) | Compound (c) | ld | lc | ldc | No. tips |
| --- | --- | --- | --- | --- | --- | --- | --- | --- |
| Geeta *et al.* [39] | 428 (76.3%) | 38 (6.8%) | 17 (3.0%) | 68 (12.1%) | 7 (1.2%) | 1 (0.2%) | 2 (0.4%) | 561 |
| Zuntini *et al.* [41] species | 2445 (86.0%) | 102 (3.6%) | 83 (2.9%) | 212 (7.5%) | - | - | - | 2842 |
| Zuntini *et al.* [41] genus | 1910 (88.3%) | 82 (3.8%) | 47 (2.2%) | 123 (5.7%) | - | - | - | 2162 |
| Janssens *et al.* [40] species | 2606 (87.2%) | 125 (4.2%) | 99 (3.3%) | 160 (5.4%) | - | - | - | 2990 |
| Janssens *et al.* [40] genus | 2178 (88.6%) | 97 (3.9%) | 55 (2.2%) | 128 (5.2%) | - | - | - | 2458 |

We used 3 different phylogenies from the literature to infer the evolutionary rates. First, to compare our predictions directly to the Geeta *et al*. [39] study, we used the same phylogeny as in their analysis [65], which is based on 3 genes: nuclear 18S rDNA and the plastid genes *rbcL* and *atpB*. Second, we used a tree by Janssens *et al*. [40], inferred using the plastid genes *rbcL* and *matK* (including *trnK*) and calibrated with 56 fossils. This tree comprises 36,101 species, representing 8,399 genera, 426 families and all orders of flowering plants (available in Supplementary material 5 of Janssens *et al*. [40]). Finally, we used a recent phylogeny by Zuntini *et al*. [41], containing 9,506 species (7,923 genera and all angiosperm families and orders), inferred using 353 nuclear genes and calibrated with 200 fossil records (available at https://zenodo.org/records/10778207; we used the tree 4-young-tree-smoothing-10-pruned-for-diversification-analyses). We show results from the Geeta *et al*. [39] phylogeny, labelled with their own classification data and the Janssens *et al*. [40] and Zuntini *et al*. [41] trees labelled with our own classification data from Naturalis Biodiversity Center [34].

Rates were inferred using the MultiState method from BayesTraits V4.1.2 [42]. We used MCMC for 1,010,000 iterations with a sample period of 1000 and removed the first 10,000 iterations to obtain posterior distributions. We used a uniform prior between 0 and 0.1 for the Zuntini *et al*. [41] trees (figs. S17 and S18) and Janssens *et al*. [40] trees (figs. S19 and S20). However, for the Geeta *et al*. [39] tree, because branch lengths are measured in substitutions per site rather than millions of years, we instead used a uniform prior between 0 and 100 (fig. S21). We took the mean of each posterior as the transition rate estimate. In addition, we estimated the rates using MLE for comparison (figs. 7a and 7b). As the units of time differ between our phylogenies and the simulations, we needed to normalise the rates to allow for comparison. We did this using the same method as for the simulations (eq. (6)): by dividing each transition rate estimate by the mean transition rate estimate for each tree.

## ACKNOWLEDGMENTS

JSM was supported by funding from the Biotechnology and Biological Sciences Research Council (UKRI-BBSRC) [grant number BB/T008784/1]. LMD was supported, in part, by National Science Foundation OISE 2020577 and a Stony Brook University Presidential Innovation and Excellence. SvdD and NSM were supported by the Issachar fund. NSM acknowledges support of the Spanish Ministry of Science and Innovation through the Centro de Excelencia Severo Ochoa (CEX2020-001049-S, MCIN/AEI/10.13039/501100011033), the EMBL partnership and the Generalitat de Catalunya through the CERCA programme. Research for this publication has been partially carried out in the Barcelona Collaboratorium for Modelling and Predictive Biology. This research is part of grant JDC2022-049526-I to NSM, funded by MCIU/AEI/10.13039/501100011033 and by European Union NextGenerationEU/PRTR.

## V. EXTENDED DATA

