## Supplementary Information for "Developmental bias explains the evolutionary trend towards simple leaf shapes"

#### CONTENTS

S1. Supplementary Simulation Methods 1
A. Leaf model details 1
1. Initialisation 1
2. Margin webbing 2
3. Margin patterning 4
4. Vein patterning 4
5. Global growth 5
6. Miscellaneous 6
B. Initial leaves 7
C. All random walk algorithms 8
D. Parameters included in the search 9
E. Agreement between manual and automatic leaf shape classification 10
F. Examples of rejected leaves 11

S2. Supplementary Phylogenetic Methods 12
A. Phylogenetic rate inference 12
B. Phylogenetic trees 13
C. Phylogenetic tree statistics 18

S3. Supplementary Results 19
A. MCMC traces 19
B. PCA occupancy for higher dimensions 23
C. Detailed balance analysis 24

S4. A short literature review of the adaptive significance of leaf shape 25
A. Introduction 25
B. Size 25
C. Teeth 25
D. Dissection and compoundness 26
1. Thermoregulation 27
2. Hydraulic efficiency 27
3. Biomechanical stability 28
4. Herbivory defence 28
5. Light interception and optimisation 28
6. Nutrient cycling 29
7. Conclusion 29

References 29

### S1. SUPPLEMENTARY SIMULATION METHODS

#### A. Leaf model details

In this section we provide a more detailed overview of how model parameters relate to growth processes. For further model details, please also see the main text and supplementary information of Runions *et al.* [1].

##### 1. Initialisation

The initial condition of the Runions *et al.* [1] simulation is a primordium (fig. S1) with user-defined dimensions and morphogen distribution along the margin. These properties are specified by the following initialisation parameters.

- NSAMP – primordium length in number of sample points.
- I_SAMPDIST – vertical distance between sample points.

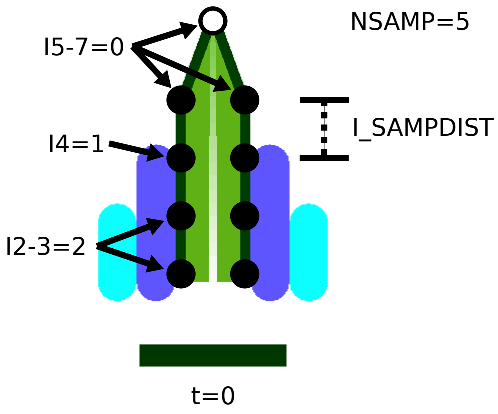


FIG. S1. **The initial leaf "primordium" at the start of the simulation labelled with a selection of initialisation parameters.** Parameters I0–I12 specify which compounds/morphogens are present at each sample point, in this case either compound 0, 1 or 2. I_SAMPDIST specifies the vertical distance between sample points and NSAMP gives the total primordium length in number of sample points.

- INITSTATE – an array containing I0–I12. Specifies which compounds are present at each sample point in the primordium.
- PRIMSCALEX – the primordium is scaled in x according to this value.
- PRIMSCALEY – the primordium is scaled in y according to this value.

##### *2. Margin webbing*

The margin parameters control patterning, webbing and tip growth. Patterning is the addition of new vein tips (convergence points) and the updating of compound positions. Webbing is the rate at which the margin deforms to reduce stretching and bending. Tip growth is the rate of extension of vein tips.

Webbing (fig. S2) is simulated for each sample point $\boldsymbol{P_i}$ along the leaf margin that is not a vein tip/convergence point, where $i$ is the sample point index. It is implemented as a displacement vector $\Delta\boldsymbol{W_i}$ that is calculated as the weighted sum of 4 other displacement vectors: stretch ($\Delta\boldsymbol{S_i}$), curvature 1 ($\Delta\boldsymbol{K_i^{(1)}}$), curvature 2 ($\Delta\boldsymbol{K_i^{(2)}}$) and normal displacement ($\boldsymbol{N_i}$).

$$\Delta\boldsymbol{W_i} = \alpha_S \Delta\boldsymbol{S_i} + \alpha_{K^{(1)}}\Delta\boldsymbol{K_i^{(1)}} + \alpha_{K^{(2)}}\Delta\boldsymbol{K_i^{(2)}} + \alpha_N \boldsymbol{N_i} \tag{1}$$

Each of these vectors is weighted by user-controlled parameters $\alpha_S$, $\alpha_{K^{(1)}}$, $\alpha_{K^{(2)}}$ and $\alpha_N$. The first three vectors ($\Delta\boldsymbol{S_i}$, $\Delta\boldsymbol{K_i^{(1)}}$, $\Delta\boldsymbol{K_i^{(2)}}$) are calculated by minimising an energy term for all sample points in the margin $\boldsymbol{P_0}, \boldsymbol{P_1}, ..., \boldsymbol{P_n}$.

The stretch energy $E_S$ is given by

$$E_S = \sum_{i=1}^{n} ||\boldsymbol{P_i} - \boldsymbol{P_{i-1}}||^2. \tag{2}$$

Curvature is approximated using two different expressions for additional flexibility. The first curvature energy is approximated for each sample point by the turning angle $\theta_i$. So for the entire margin this results in the curvature energy

$$E_{K^{(1)}} = \sum_{i=1}^{n-1} \theta_i^2. \tag{3}$$

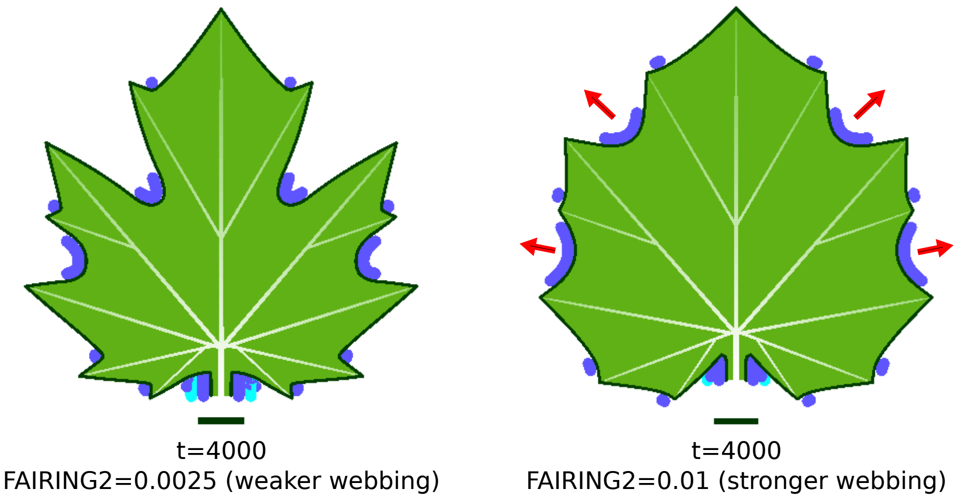


FIG. S2. **Stronger webbing results in shallower lobes.** Stronger webbing results in faster growth of the margin and lamina between vein tips (convergence points) to reduce stretching. The overall rate of webbing depends on multiple morphogen-associated model parameters. In this example, compound 2 (dark blue) has stronger webbing for the right-hand leaf, resulting in faster growth in the lobes where it is localised.

The second curvature energy is given by the weighted second finite difference about each sample point

$$E_{K^{(2)}} = \sum_{i=1}^{n-1} ||w_l^i \boldsymbol{P_{i-1}} - w_c^i \boldsymbol{P_i} + w_r^i \boldsymbol{P_{i+1}}||^2, \tag{4}$$

where $w_l^i = w_r^i = \frac{1}{2}$ and $w_c^i = 1$. These weights result in a vector from $\boldsymbol{P_i}$ to the mid-point of the line between $\boldsymbol{P_{i-1}}$ and $\boldsymbol{P_{i+1}}$.

Finally the magnitude of the normal displacement vector for each sample point $\boldsymbol{P_i}$ is proportional to the average length of the adjacent margin edges

$$\frac{1}{2}\big(||\boldsymbol{P_{i+1}} - \boldsymbol{P_i}|| + ||\boldsymbol{P_i} - \boldsymbol{P_{i-1}}||\big). \tag{5}$$

To calculate the overall webbing vector for each sample point, each of these terms is weighted by a user-controlled parameter: stretch ($\alpha_S$) - STRETCH, curvature 1 ($\alpha_{K^{(1)}}$) - CFLOW, curvature 2 ($\alpha_{K^{(2)}}$) - CURVATURE, normal displacement ($\alpha_N$) - NORMAL.

Many margin parameters are associated with particular morphogens/compounds and have the name format <parameter_name><morphogen_number>, e.g. FAIRING2. For a particular margin segment, not all margin parameters are necessarily used, as the parameters used depend on the compounds present there. If there are 2 compounds present, each will have its own set of parameters actively playing a role in growth and form. Webbing can be controlled by the user via the following parameters.

- TIP_GROWTH – controls the rate at which vein tips are displaced in the vein direction.
- FAIRING – controls the total rate of webbing.
- CURVATURE – controls the rate of bending.
- CFLOW – controls the rate of bending.
- STRETCH – controls the distance between neighbouring margin points.
- NORMAL – the amount that margin points are displaced normal to the curve.

##### *3. Margin patterning*

The margin is patterned by convergence points, which mark the tips of new veins, and compounds (also referred to as morphogens) (fig. S3). The patterning process occurs by first measuring distances along the margin. This informs where convergence points and thus veins should be added. The margin points at which compounds are expressed are updated in response to the new convergence points. Finally, new veins are added to connect the convergence points to the existing vasculature.

The following parameters control margin patterning globally.

- LASSYMETRY – the distance value on the first (left) point along the margin. Can be thought of as the effect of neighbouring tissue on the leaf primordium.
- RASSYMETRY – the distance value on the last (right) point along the margin.
- INTERVAL – the distance along the margin between two convergence points over which a compound will be expressed.

The following parameters have compound-specific effects.

- METRIC – the rate at which distances along the margin increase relative to arc-length distances. Measured distances can increase more slowly or more rapidly than the arc-length distance as the leaf grows depending on the value here.
- CP_DIST – new convergence points are added to a margin segment when the distance measured along this segment exceeds this value. This may differ between compounds.
- PATTERNABLE – controls whether margin segments expressing this compound are competent for new convergence points.
- COMPETENCE – sets the maximum distance from the leaf base a margin segment can be and still be competent for new convergence points.
- COMPETENCET – maximum time step beyond which a margin segment will no longer be competent for new convergence points.
- EXTENDABLE – whether the interval in which a compound is expressed can increase in size or not.
- EXTENDABLE_CONTEXT – which compounds can be overwritten. When a compound number is greater than this value, it can be overwritten.

##### *4. Vein patterning*

New veins attach convergence points to existing vasculature so as to minimise resistance (fig. S4). Resistance depends on the distance travelled in the blade and the distance travelled in the vasculature to the leaf base. The user can exert some control over vein attachment through the following parameters.

- AXISANGLE – the angle at which new veins branch from existing veins.
- MAXANGLE – the maximum angle veins can make with the margin. This restricts the orientation of new veins.
- GLOBAL_RESISTANCE – determines how the resistance (length) of veins affects where a vein will join to the existing vasculature. A value of 0 means only the resistance of the new vein segment is considered. A value of 1 means the total resistance of the entire path to the leaf base is considered.

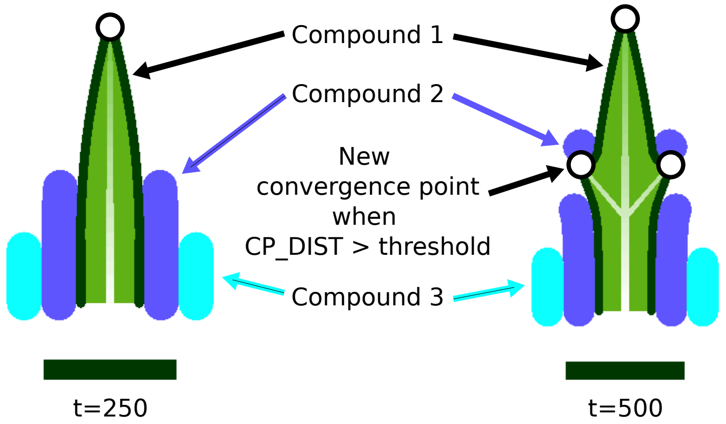


FIG. S3. **Margins are patterned by convergence points and morphogens/compounds.** New convergence points are inserted between existing convergence points, where the intervening distance along the margin exceeds a threshold value. Compounds/morphogens on the margin can alter the measured distances between convergence points, affecting where new convergence points are inserted. New veins are inserted to connect new convergence points to the existing vasculature, thereby specifying new growth axes. Compounds can also alter the rate of webbing between veins.

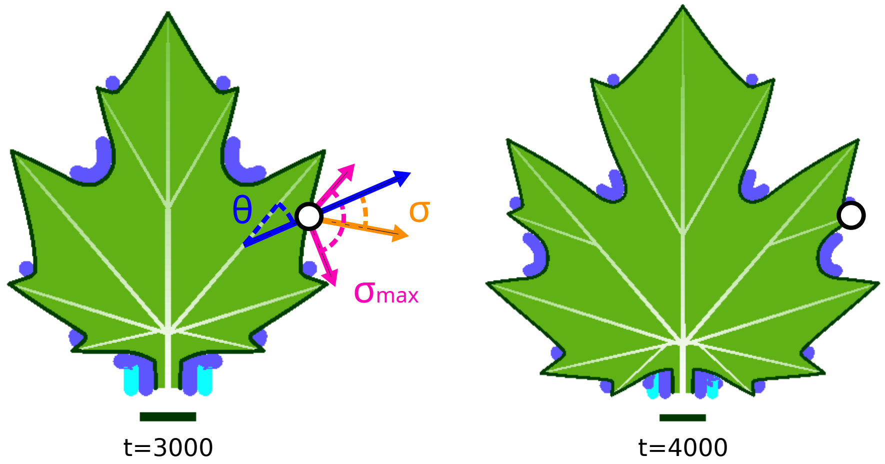


FIG. S4. **New veins attach to existing vasculature at angle $\theta$.** Attachment is restricted such that the angle made with the margin $\sigma$ cannot exceed $\sigma_{\max}$. Veins attach so as to minimise "resistance"—the total distance from the vein tip to either the base of the leaf or the existing vasculature, depending on the value of GLOBAL_RESISTANCE.

##### *5. Global growth*

Growth is a combination of uniform scaling, vein tip displacement, webbing and vein extension. The global rate of tip displacement, webbing and vein extension depends on the distance to the leaf base as measured along the veins according to a growth gradient (fig. S5). Vein growth rate can also be age-dependent.

- UNIFORM_GROWTH – the rate of uniform scaling.
- TG_START_X – the distance from the leaf base along the veins that the rate of tip growth begins to decrease.
- TG_END_X – the distance from the leaf base along the veins that the rate of tip growth reaches its minimum.
- FG_START_X – the distance from the leaf base that the rate of webbing growth begins to fall off.
- FG_END_X – the distance from the leaf base that the rate of webbing growth stops decreasing.

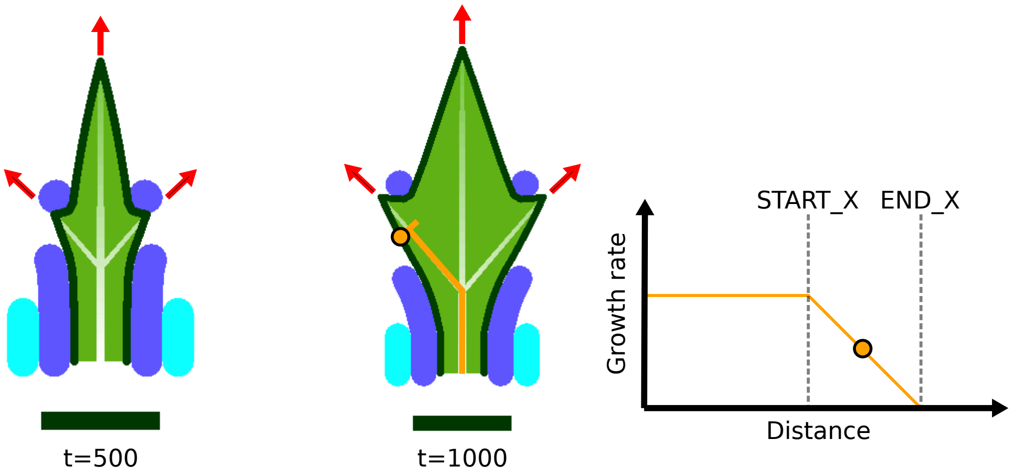


FIG. S5. **Veins are the main axes of growth, pushing the local margin outwards in successive time points**. The rate of growth of the veins and margin depends on the distance to the base of the leaf. Growth of veins and margin is fastest near the base of the leaf, tailing off towards the tip depending on the values of parameters such as VG_START_X for vein growth.

- VG_GRATE – base vein growth rate.
- VG_START_X – distance from leaf base that the rate of vein growth begins to fall off.
- VG_END_X – distance from leaf base that the rate of vein growth stops decreasing.
- CP_AD_TIP_GROWTH – rate of extra early tip growth.
- CP_ADTG_FALLOF – time period over which early growth disappears.
- AX_AD_AXIS_GROWTH – rate of extra early vein-extension.
- AX_ADAG_FALLOF – time period over which early vein extension growth disappears.

##### *6. Miscellaneous*

- APEX_TIP_GROWTH – extra tip growth for the primary axis.
- ASYMMETRIC_FAIRING – whether the rate of webbing can be different on either side of a convergence point.
- ASSYM_PRE – the rate of webbing before the convergence point.
- ASSYM_POST – the rate of webbing after the convergence point.
- SHARP_LEAF_TIP – whether to disable webbing at the leaf tip.
- INTERNAL_FLOW – whether webbing can only occur in the outward direction.
- PETIOLE_COMPOUND – the compound which specifies the petiole.
- PETIOLE_EXTEND – whether the petiole (as marked by PETIOLE_COMPOUND) is enlarged by uniform growth.

##### B. Initial leaves

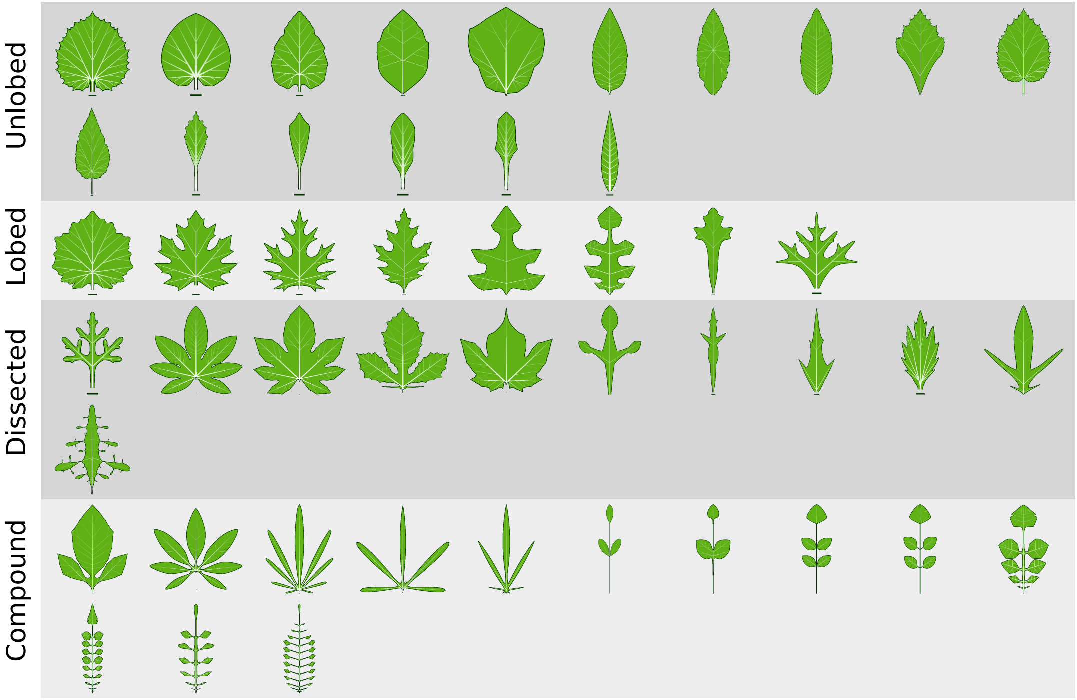


FIG. S6. **The sample of 48 leaves used as the starting points for random walks.** 16 unlobed (u), 8 lobed (l), 11 dissected (d) and 13 compound (c).

##### C. All random walk algorithms

**Algorithm S1:** Random Walk MUT2

```
θ ← initial parameter values
for step ← 0 to N_maxGenerated − 1 do
    for attempt ← 0 to N_maxAttempts − 1 do
        θ′ ← θ                                          // Copy parameters for this step
        mutation ← random draw from {
            uniform[0.1, 1),
            uniform[1, 10),
            uniform[10, 100)
        }
        sign ← random choice from {−1, 1}
        i ← random draw from {0, . . . , length(θ) − 1}
        θ′[i] ← θ[i] + (mutation × sign)                // Mutate
        Run simulation with θ′
        if leaf passes sanity checks then
            θ ← θ′                                      // Save new parameters
            break
```

**Algorithm S2:** Random Walk MUT5

```
θ ← initial parameter values
for step ← 0 to N_maxGenerated − 1 do
    for attempt ← 0 to N_maxAttempts − 1 do
        θ′ ← θ                                          // Copy parameters for this step
        mutation ← random draw from {
            uniform[0.1, 1),
            uniform[1, 10),
            uniform[10, 100)
        }
        sign ← random choice from {−1, 1}
        scale ← 0.1 × initialSampleRange    // This is the range within the leaves in Figure S6
        i ← random draw from {0, . . . , length(θ) − 1}
        θ′[i] ← θ[i] + (mutation × sign × scale)        // Mutate
        Run simulation with θ′
        if leaf passes sanity checks then
            θ ← θ′                                      // Save new parameters
            break
```

##### D. Parameters included in the search

| Parameter | Target | Parameter | Target | Parameter | Target |
|---|---|---|---|---|---|
| PSPACE1 | 1 | FAIRING3 | 1 | CP_DIST | 1 |
| PSPACE2 | 1 | CURVATURE3 | 1 | FILLNONAPICALINTERVALS | 1 |
| DISPSTEP | 1 | STRETCH3 | 1 | NCOMP | 0 |
| GDT | 1 | CFLOW3 | 1 | INTERVAL1 | 1 |
| FINALFRAME | 1 | NORMAL3 | 1 | INTERVAL2 | 1 |
| NUMFLOW | 1 | TIP_GROWTH3 | 1 | INTERVAL3 | 1 |
| NSAMP | 1 | CPDIST3 | 1 | INTERVAL4 | 1 |
| I_SAMPDIST | 1 | METRIC3 | 1 | NEWAXISINTS | 1 |
| I0 | 1 | EXTENDABLE3 | 1 | NEWAXISINTE | 1 |
| I1 | 1 | EXTENDABLE_CONTEXT3 | 1 | FAIRING1 | 1 |
| I2 | 1 | PATTERNABLE3 | 1 | CURVATURE1 | 1 |
| I3 | 1 | COMPETENCE3 | 1 | STRETCH1 | 1 |
| I4 | 1 | COMPETENCET3 | 1 | CFLOW1 | 1 |
| I5 | 1 | FAIRING4 | 1 | NORMAL1 | 1 |
| I6 | 0 | CURVATURE4 | 1 | TIP_GROWTH1 | 1 |
| I7 | 1 | STRETCH4 | 1 | CPDIST1 | 1 |
| I8 | 1 | CFLOW4 | 1 | METRIC1 | 1 |
| I9 | 1 | NORMAL4 | 1 | EXTENDABLE1 | 1 |
| I10 | 1 | TIP_GROWTH4 | 1 | EXTENDABLE_CONTEXT1 | 1 |
| I11 | 1 | CPDIST4 | 1 | PATTERNABLE1 | 1 |
| I12 | 1 | METRIC4 | 1 | COMPETENCE1 | 1 |
| PRIMSCALEX | 1 | EXTENDABLE4 | 1 | COMPETENCET1 | 1 |
| PRIMSCALEY | 1 | EXTENDABLE_CONTEXT4 | 1 | FAIRING2 | 1 |
| LASSYMETRY | 1 | PATTERNABLE4 | 1 | CURVATURE2 | 1 |
| RASSYMETRY | 1 | COMPETENCE4 | 1 | STRETCH2 | 1 |
| DIVTHRESH | 1 | COMPETENCET4 | 1 | CFLOW2 | 1 |
| MAXANGLE | 1 | AXISANGLE | 1 | NORMAL2 | 1 |
| TIP_GROWTH2 | 1 | CPDIST2 | 1 | METRIC2 | 1 |
| EXTENDABLE2 | 1 | EXTENDABLE_CONTEXT2 | 1 | PATTERNABLE2 | 1 |
| COMPETENCE2 | 1 | COMPETENCET2 | 1 | VG_START_X | 0 |
| VG_END_X | 1 | CP_AD_TIP_GROWTH | 1 | CP_ADTG_FALLOFF | 1 |
| AX_AD_AXIS_GROWTH | 0 | AX_ADAG_FALLOFF | 0 | APEX_TIP_GROWTH | 1 |
| ASYMMETRIC_FAIRING | 1 | ASYMM_PRE | 0 | ASYMM_POST | 0 |
| SHARP_LEAF_TIP | 1 | INTERNAL_FLOW | 1 | PETIOLE_COMPOUND | 0 |
| PETIOLE_EXTEND | 0 | QUADD_ERR_THRESH | 0 | COMPWIDTH | 0 |
| BB_WIDTH_MULT | 0 | MARGIN_WIDTH | 1 | SCALE_WIDTH_BY_BB | 0 |
| SUPRESSED_COMP | 0 | PM_POW | 0 | PM_WIDTH | 0 |
| PM_INC | 0 | PM_SAMP | 0 | DEFINE_TERMINAL_LEAFLET_BASE | 0 |
| BOUNDARY_MORPHOGEN | 0 | TG_START_X | 0 | TG_END_X | 1 |
| FG_START_X | 0 | FG_END_X | 1 | VG_GRATE | 1 |

TABLE S1. **Leaf developmental model [1] parameter names and whether they are allowed to vary in the random walk search (split across three columns).** A value of 1 in the Target column means the parameter was targeted in the walk, 0 means it was not. We split the INITSTATE parameter, which is a vector, up into 13 separate parameters I0–12. I6 is kept at 0; all other parameters are allowed to vary but the values of I0–5 are set to be the reverse of I7–I12. This helps promote symmetrical growth and reduce the occurrence of simulation errors and invalid leaves. Parameters were excluded from the search because some were purely graphical parameters, while others did not vary between the leaves presented in Runions *et al.* [1] and were therefore treated as not important determinants of shape. 100/122 parameters were targeted during the simulations.

##### E. Agreement between manual and automatic leaf shape classification

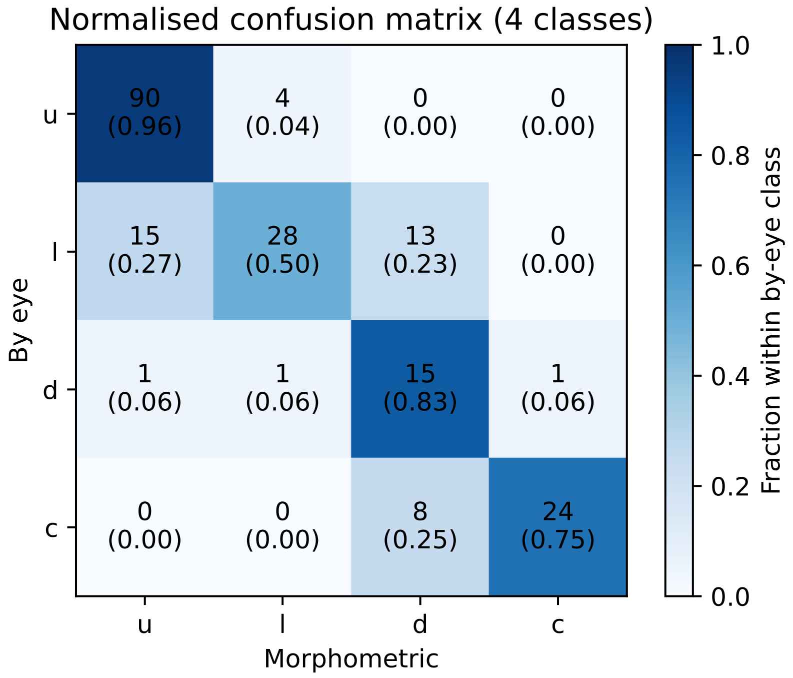


(a)

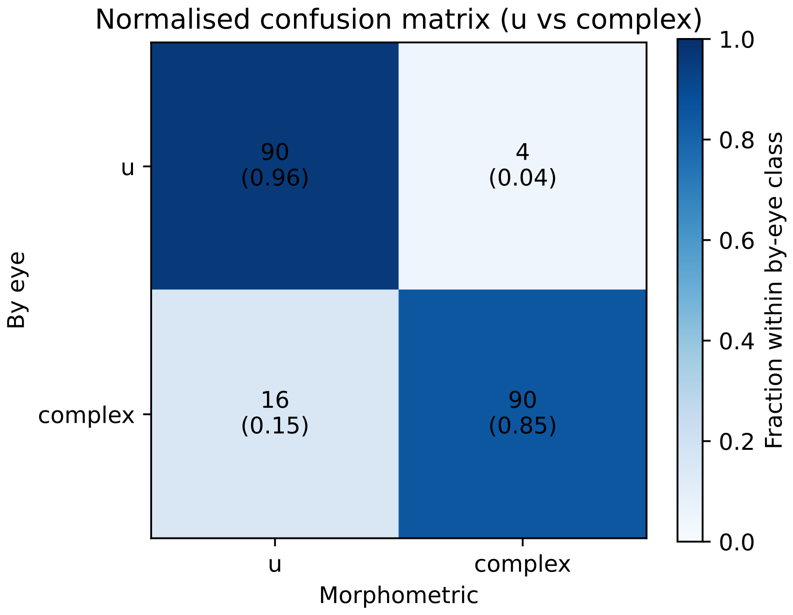


(b)

FIG. S7. **Confusion matrices comparing by-eye and automatic morphometric classification at two resolutions for a sample of 200 simulation leaf images. (a)** Four-class confusion matrix (u = unlobed, l = lobed, d = dissected, c = compound). **(b)** Binary confusion matrix after collapsing to u and complex, where complex = l ∪ d ∪ c. In both panels, cell colour shows the proportion within each by-eye class, overlaid numbers show raw counts. This comparison is also discussed in methods section IV D.

##### F. Examples of rejected leaves

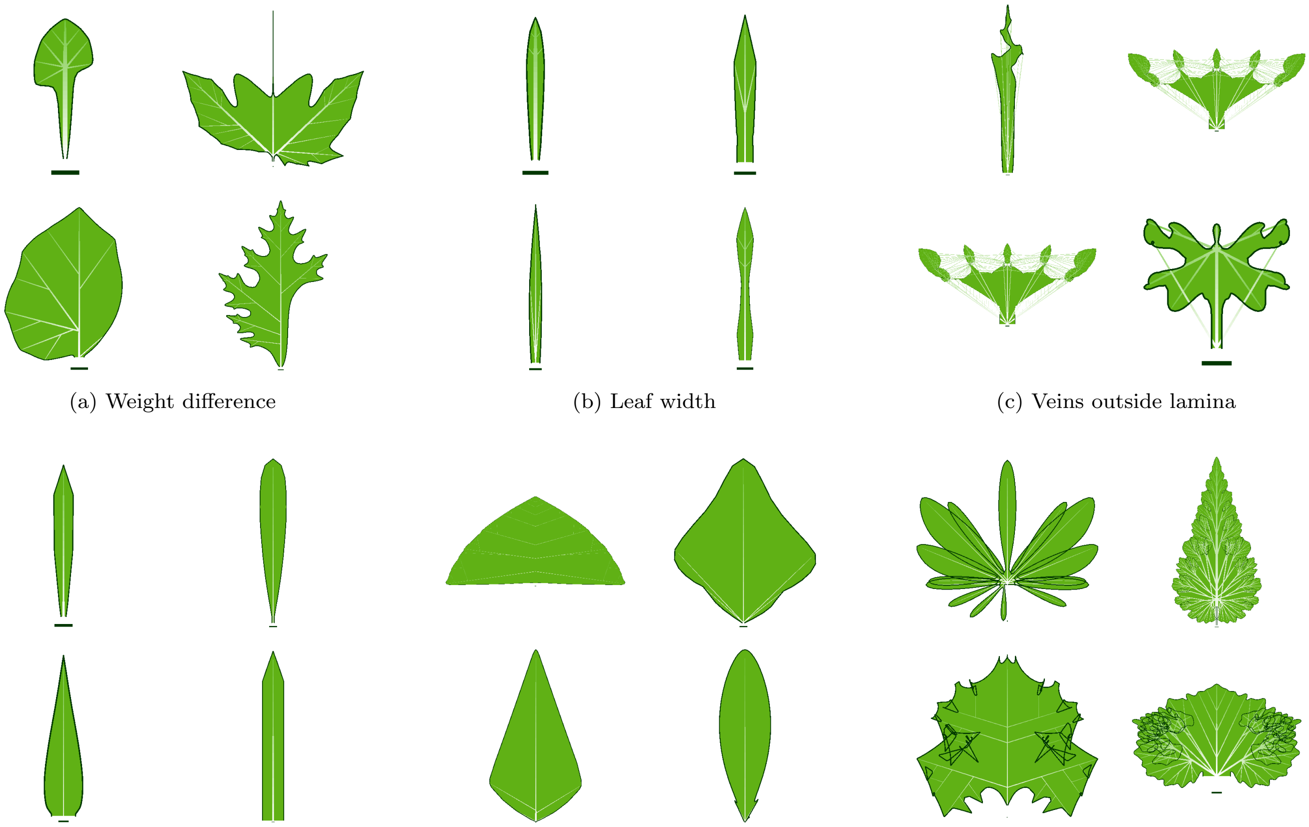

(a) Weight difference (b) Leaf width (c) Veins outside lamina

(d) Veins width (e) Veins area (f) Overlapping margin

FIG. S8. **Examples of rejected leaves with the condition (table I) that each failed to satisfy.** Each leaf shown here passed all conditions except the one stated; however, in general many leaves that are rejected fail multiple conditions.

#### S2. SUPPLEMENTARY PHYLOGENETIC METHODS

##### A. Phylogenetic rate inference

We use the MultiState method from BayesTraits V4 to infer transition rates from phylogenetic data. MultiState fits a continuous-time Markov chain (CTMC) to a discrete character distributed across the tips of a phylogeny. The tree topology and character states at the tips are fixed, with only the CTMC parameters defined by the matrix $\boldsymbol{Q}$ being estimated.

The optimal $\boldsymbol{Q}$ is found by maximising the likelihood $L$ of the observed tip data $D$ given the phylogenetic tree $T$ (topology and branch lengths) and the value of $\boldsymbol{Q}$ [2]. Let $T = (V, E)$ denote the tree, with leaf nodes $V_{\text{tip}} \subset V$ which have observed states $D_v$ for $v \in V_{\text{tip}}$, and internal nodes $V_{\text{int}} = V \setminus V_{\text{tip}}$ (including the root) which have unobserved states. Under the assumptions of the CTMC, the probability of a single branch $(u, v) \in E$ with parent state $x_u$, child state $x_v$ and branch length $t_{uv}$ is

$$p(x_u, x_v, t_{uv}) = \left[\exp(\boldsymbol{Q}t_{uv})\right]_{x_u x_v} \tag{6}$$

The likelihood is then the product of the probabilities for all branches in the tree summed over all possible assignments of character states at internal nodes [2].

$$L(D|\boldsymbol{Q}, T) = \sum_{\{x_v\}_{v \in V_{\text{int}}}} \prod_{(u,v) \in E} p(x_u, x_v, t_{uv}). \tag{7}$$

The pruning algorithm [3] is used to compute this likelihood more efficiently. The likelihood surface for $\boldsymbol{Q}$ is then searched, in the case of MLE, to find the value of $\boldsymbol{Q}$ that gives the maximum likelihood solution via the BOBYQA algorithm [4]. In the case of MCMC, a posterior distribution is sampled using Metropolis-Hastings [5]. This posterior is given by

$$p(\boldsymbol{Q}|D, T) = \frac{p(D|\boldsymbol{Q}, T)p(\boldsymbol{Q})}{\int_{\boldsymbol{Q}} p(D|\boldsymbol{Q}, T)p(\boldsymbol{Q})d\boldsymbol{Q}} \tag{8}$$

where $p(D|\boldsymbol{Q}, T) = L(D|\boldsymbol{Q}, T)$, $p(\boldsymbol{Q})$ is the prior distribution and the denominator integrates over all possible values of $\boldsymbol{Q}$. We use uniform priors between 0 and 0.1 for all trees except Geeta *et al.* [6] for which we use a uniform prior between 0 and 100. This is because branch lengths for Geeta *et al.* [6] are measured in substitutions per site rather than millions of years.

All non-diagonal elements of $\boldsymbol{Q}$ are permitted to vary freely during the fitting process. Diagonal elements are set to the negative sum of other elements in the same row, such that rows sum to zero.

Elements $q_{ij}$ of $\boldsymbol{Q}$ represent the instantaneous rate of transition between state $i$ and $j$ in units of transitions per unit of branch length. To allow comparison between phylogenies with different branch lengths, we normalise the $\boldsymbol{Q}$ matrix for each tree by dividing each transition rate by the mean of the off-diagonal elements of $\boldsymbol{Q}$ (methods section IV F).

##### B. Phylogenetic trees

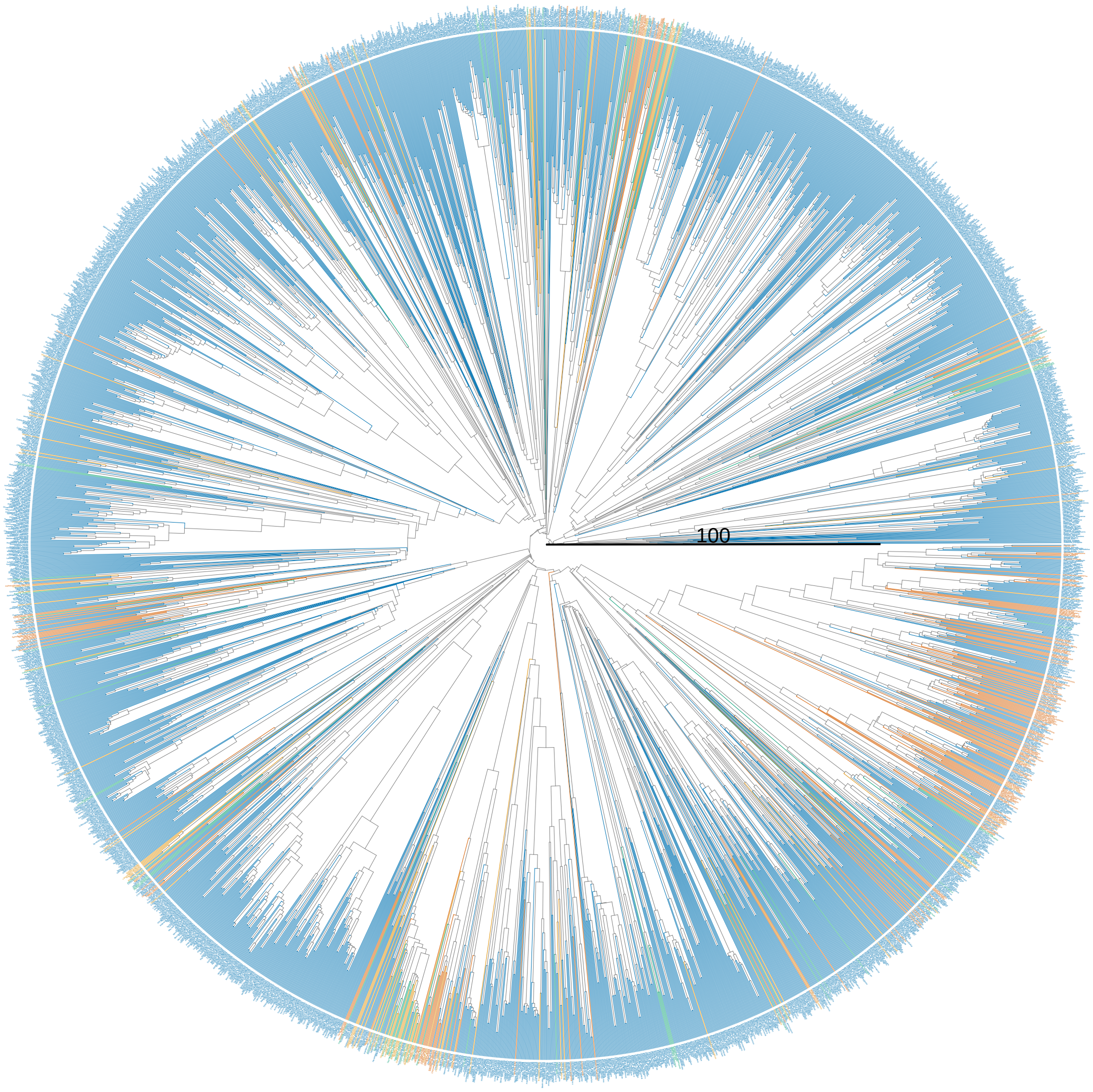


FIG. S9. Zuntini *et al.* [7] species. Branch length is millions of years (Myr).

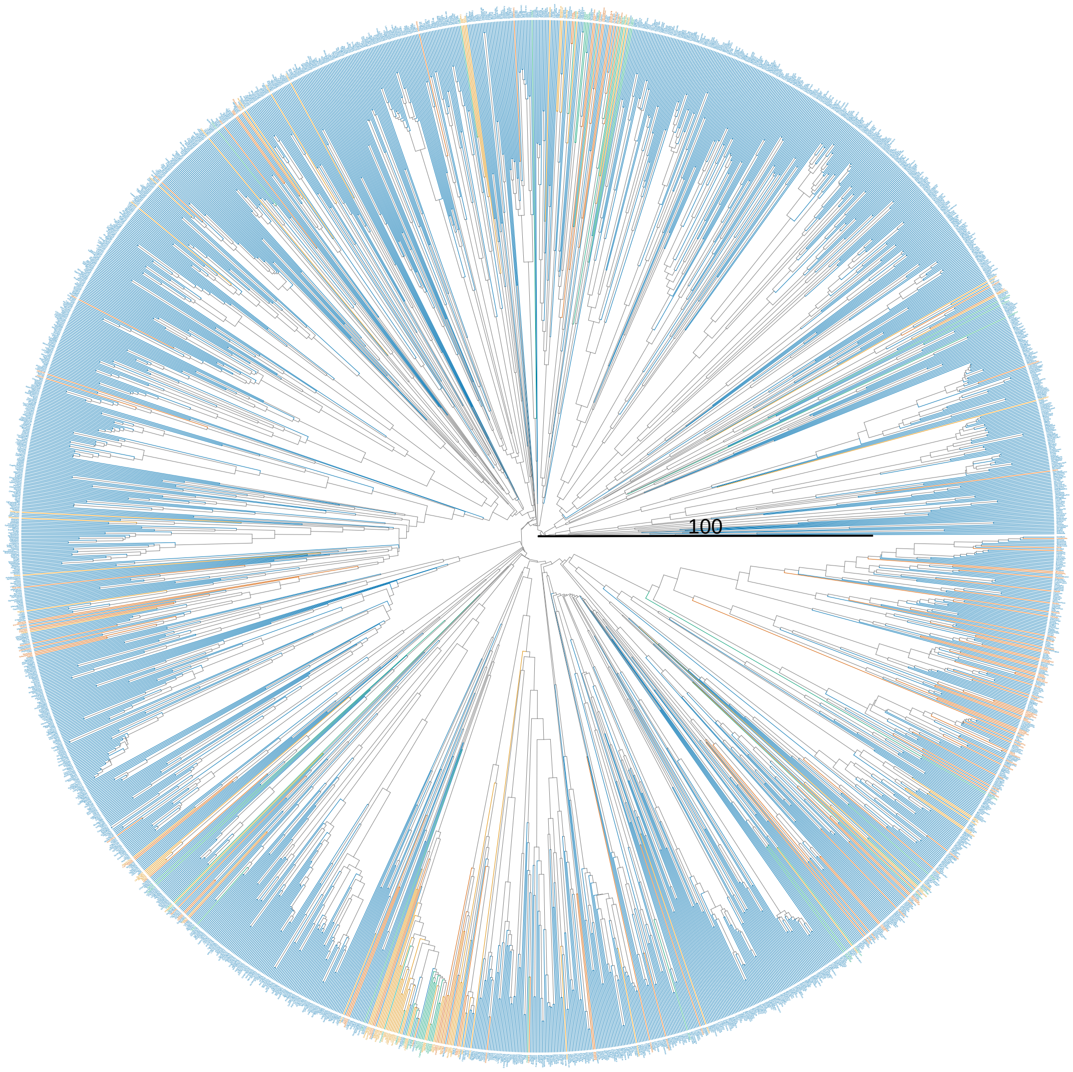


FIG. S10. Zuntini *et al.* [7] genus. Branch length is millions of years (Myr).

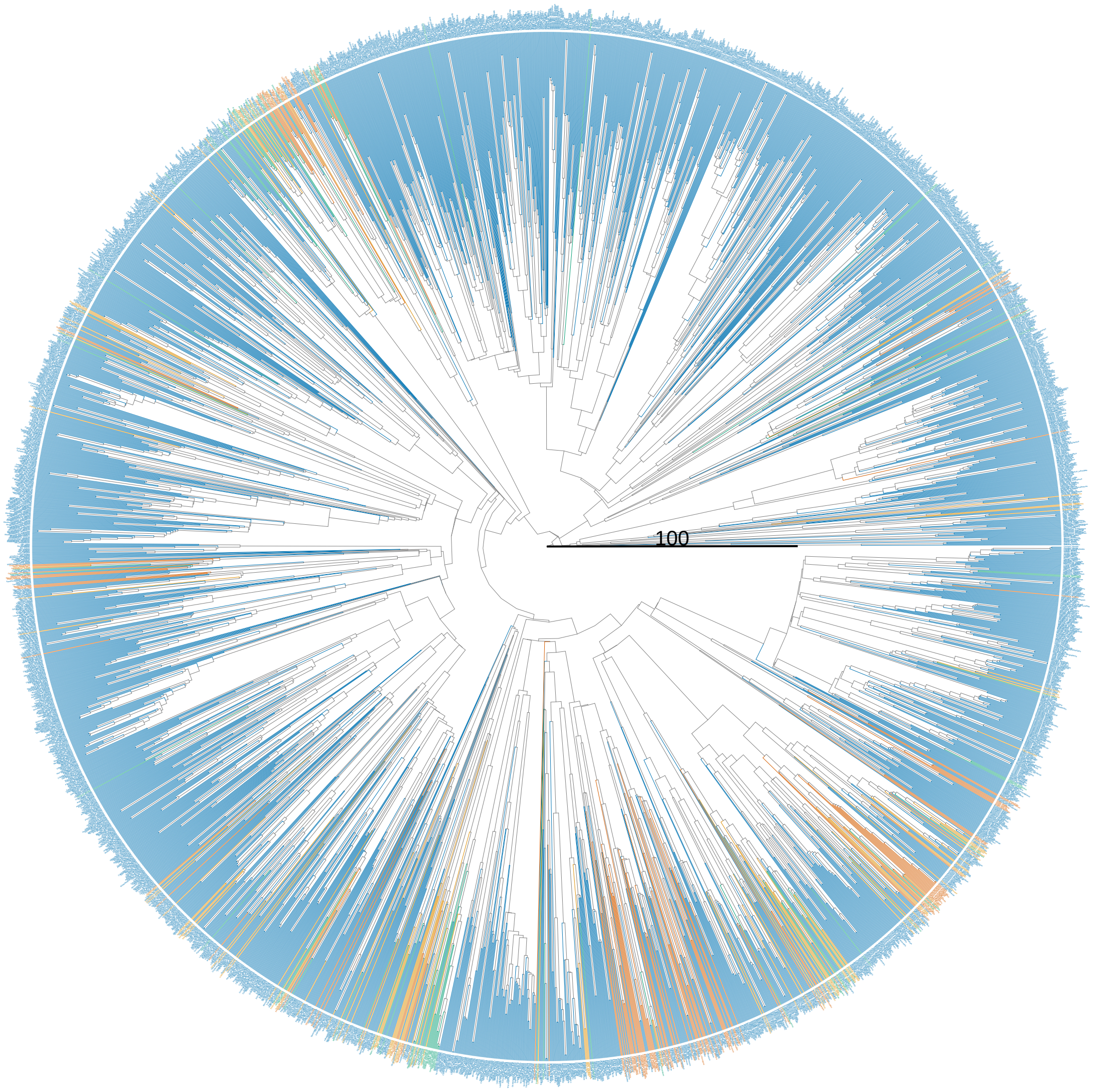


FIG. S11. Janssens *et al.* [8] species. Branch length is millions of years (Myr).

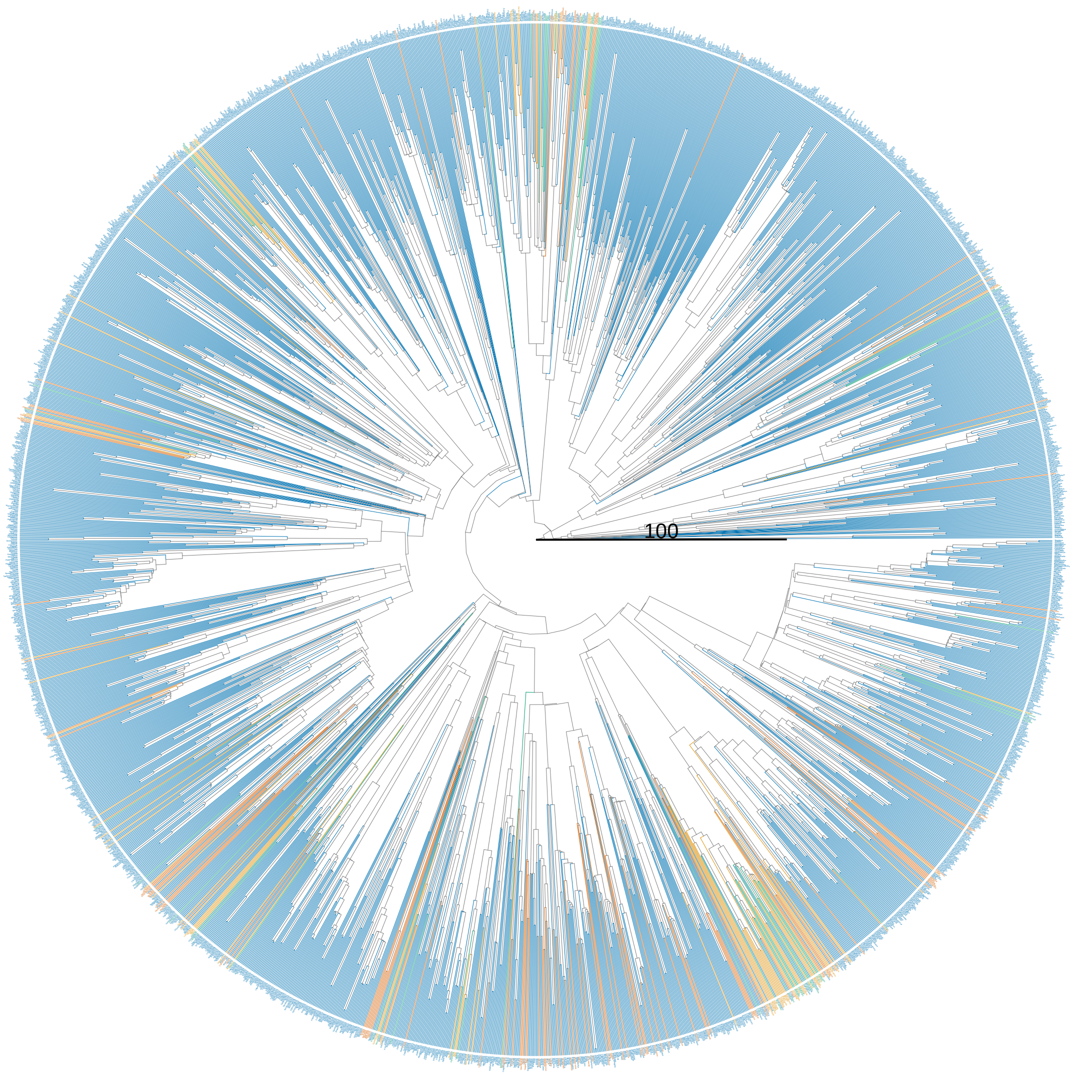


FIG. S12. Janssens *et al.* [8] genus. Branch length is millions of years (Myr).

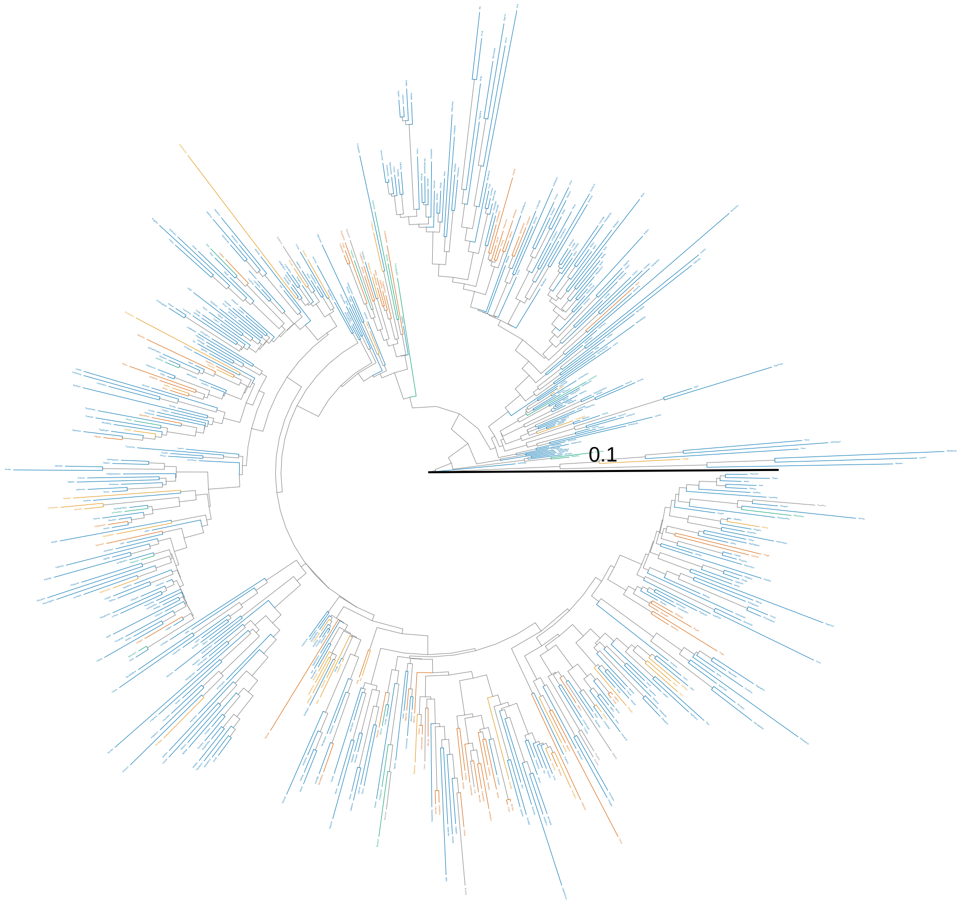


FIG. S13. Geeta *et al.* [6]. Branch length is substitutions per site.

#### C. Phylogenetic tree statistics

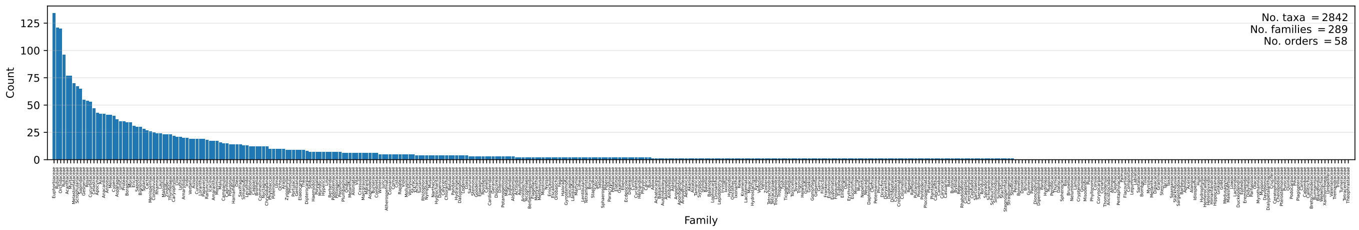


(a) Zuntini *et al.* [7] species.

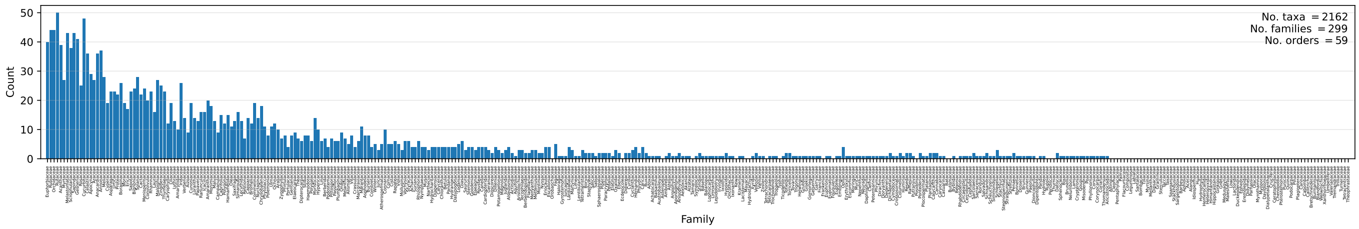


(b) Zuntini *et al.* [7] genus.

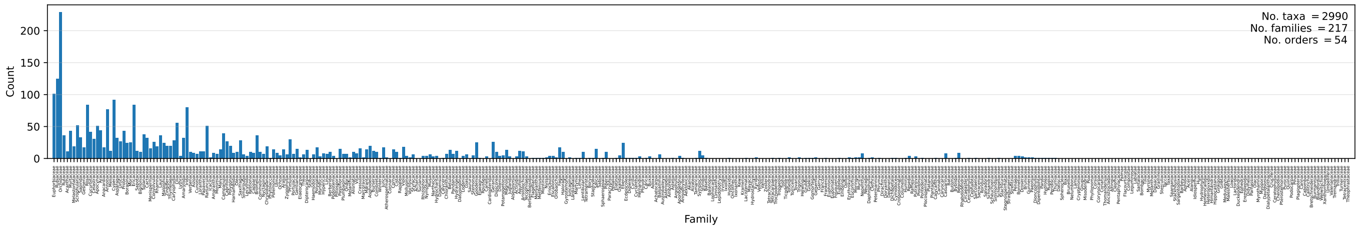


(c) Janssens *et al.* [8] species.

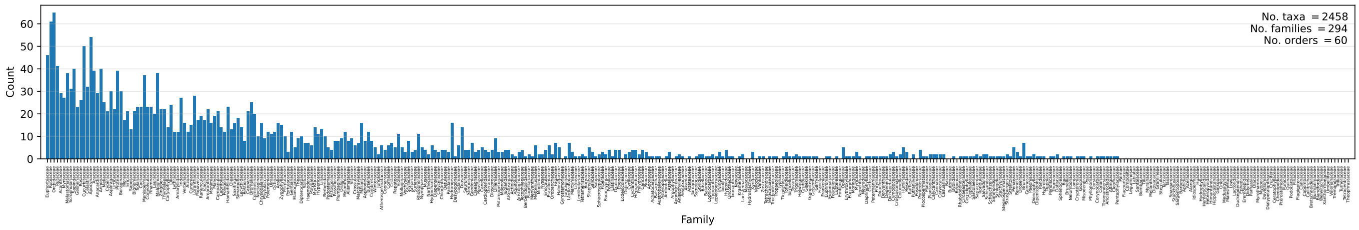


(d) Janssens *et al.* [8] genus.

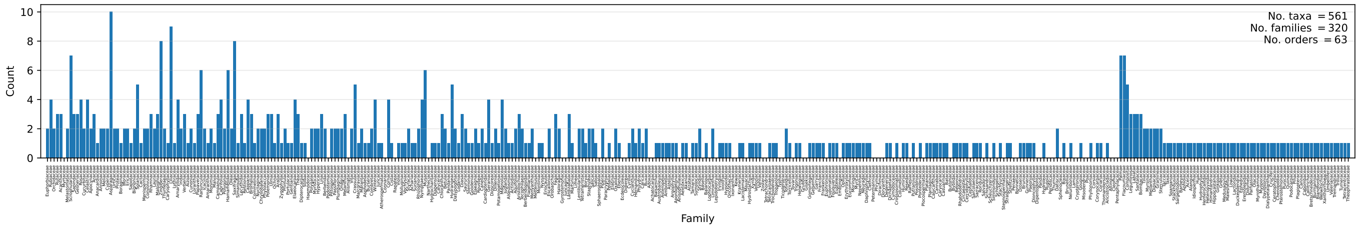


(e) Geeta *et al.* [6].

FIG. S14. **The number of taxa per family across the phylogenetic trees used in our analysis.** The distributions are more similar for the Zuntini *et al.* [7] genus and Janssens *et al.* [8] genus trees than the species trees. This could account for the increased variation in inferred shape transition rates observed between the two species trees relative to the two genus trees (figs. 7a and 7b). Families were assigned to tree tips using data from Naturalis Biodiversity Center [9].

### S3. SUPPLEMENTARY RESULTS

#### A. MCMC traces

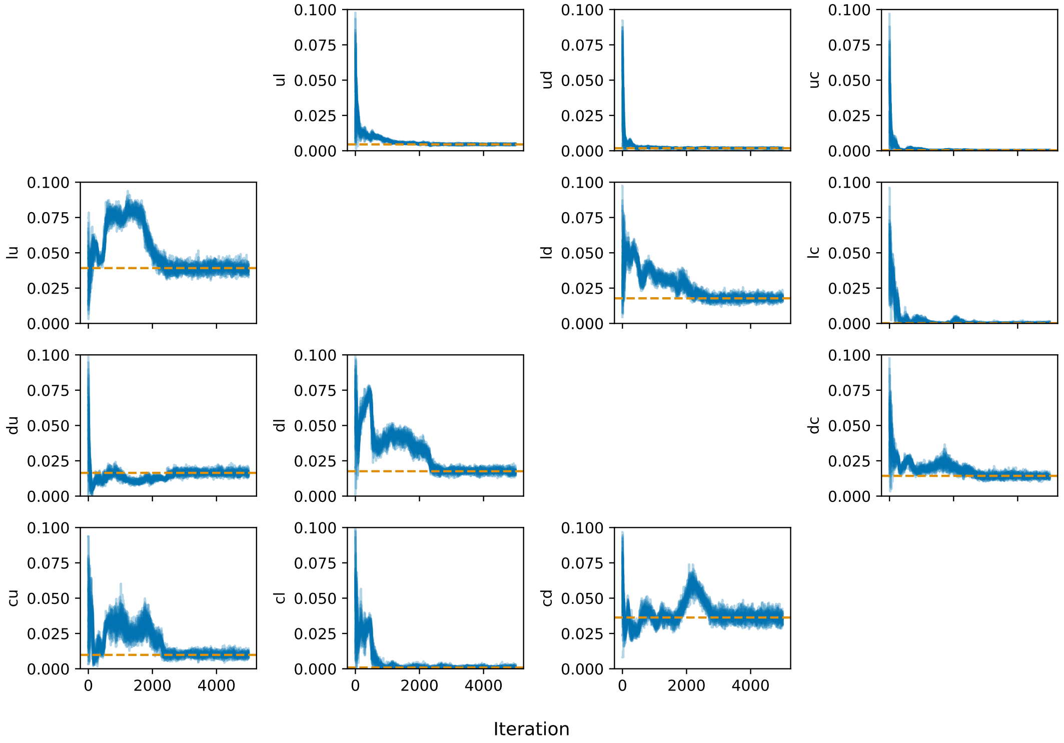


FIG. S15. **MUT2 trace.** The y-axis label gives the transition (e.g. ul = unlobed → lobed). Uniform prior between 0 and 0.1. No. chains, 24. No. iterations 5000. Burn in period, 2500 iterations. Sample period, 10 iterations. The maximum likelihood (ML) estimates for the rates are given by the orange dashed lines. Calculated using emcee package [10].

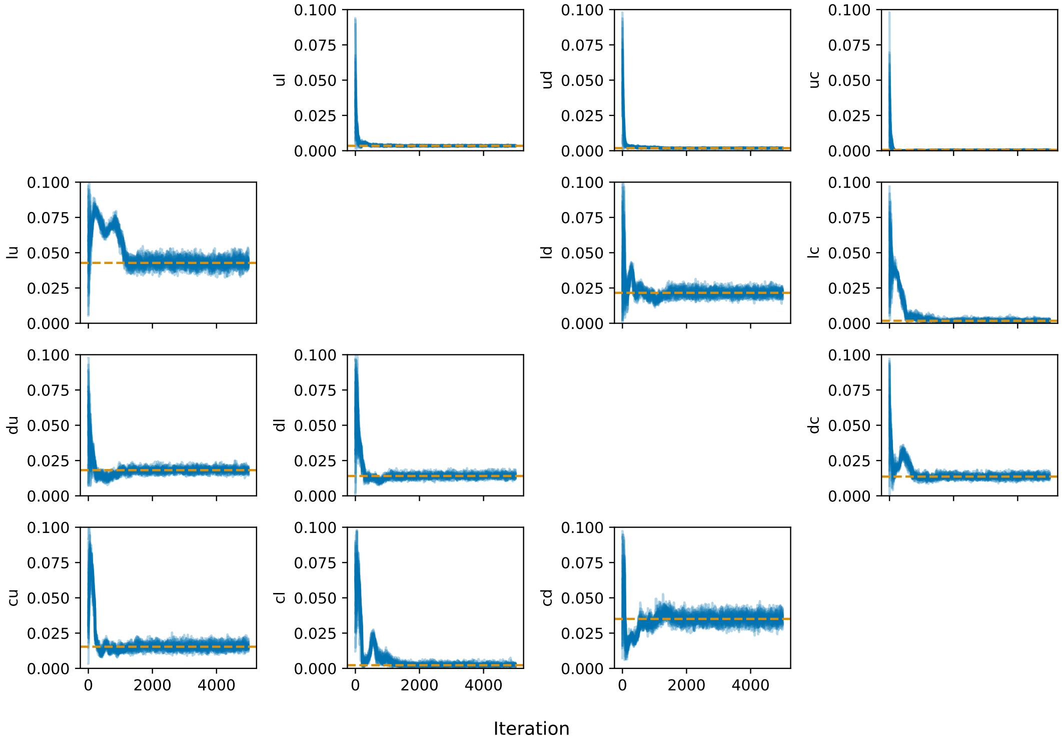


FIG. S16. **MUT5 trace.** Uniform prior between 0 and 0.1. No. chains, 24. No. iterations, 5000. Burn in period, 2500 iterations. Sample period, 10 iterations. ML estimates are given by the orange dashed lines. Calculated using emcee package [10].

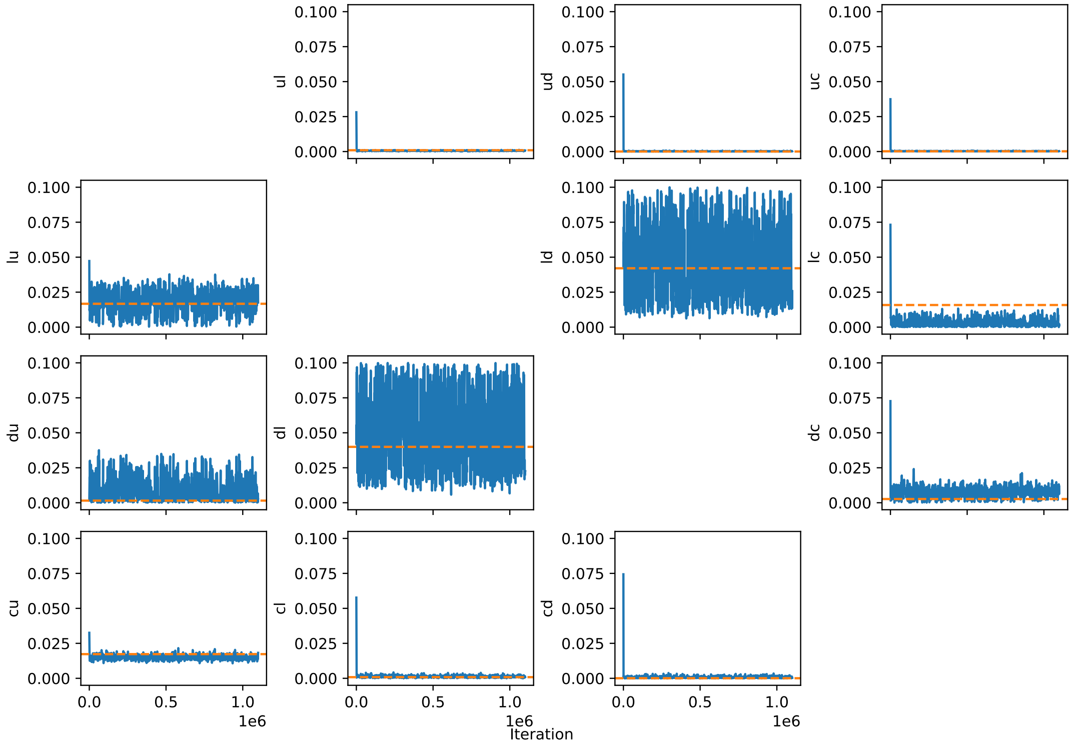


FIG. S17. **Zuntini *et al.* [7] species trace.** Uniform prior between 0 and 0.1. Total no. iterations, 1100000. Burn in period, 100000 iterations. Sample period 1000 iterations. ML estimates given by the orange dashed line. Calculated using BayesTraits V4.1.2 [11].

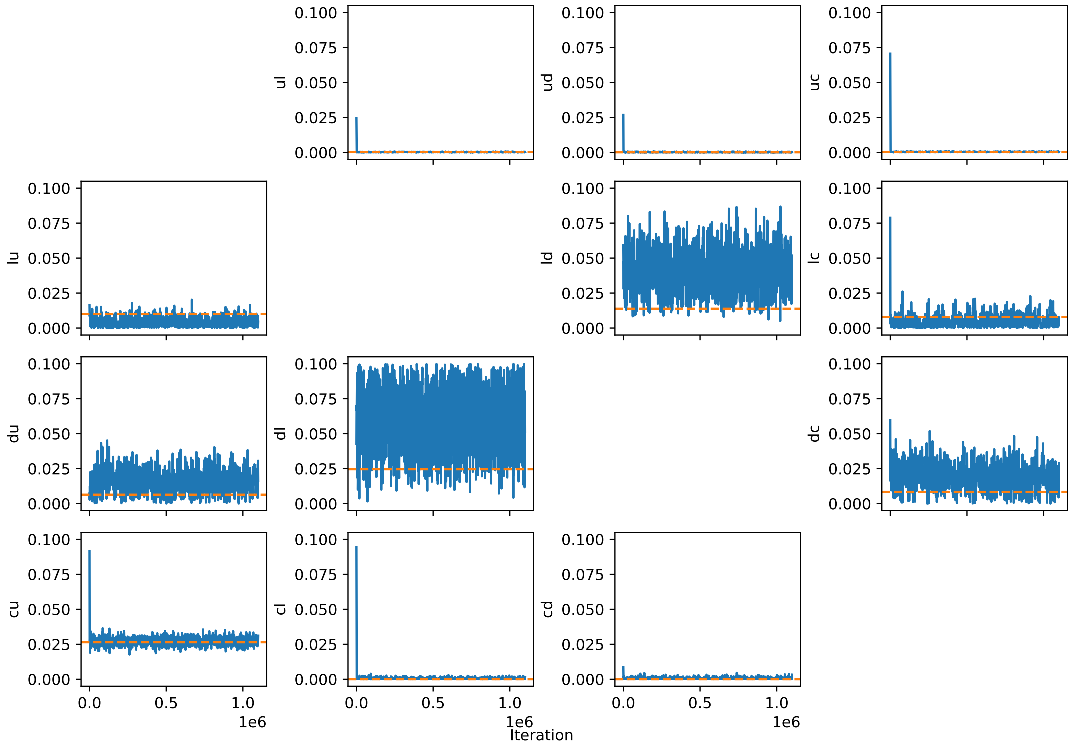


FIG. S18. **Zuntini *et al.* [7] genus trace.** Uniform prior between 0 and 0.1. Total no. iterations, 1100000. Burn in period, 100000 iterations. Sample period 1000 iterations. ML estimates given by the orange dashed line. Calculated using BayesTraits V4.1.2 [11].

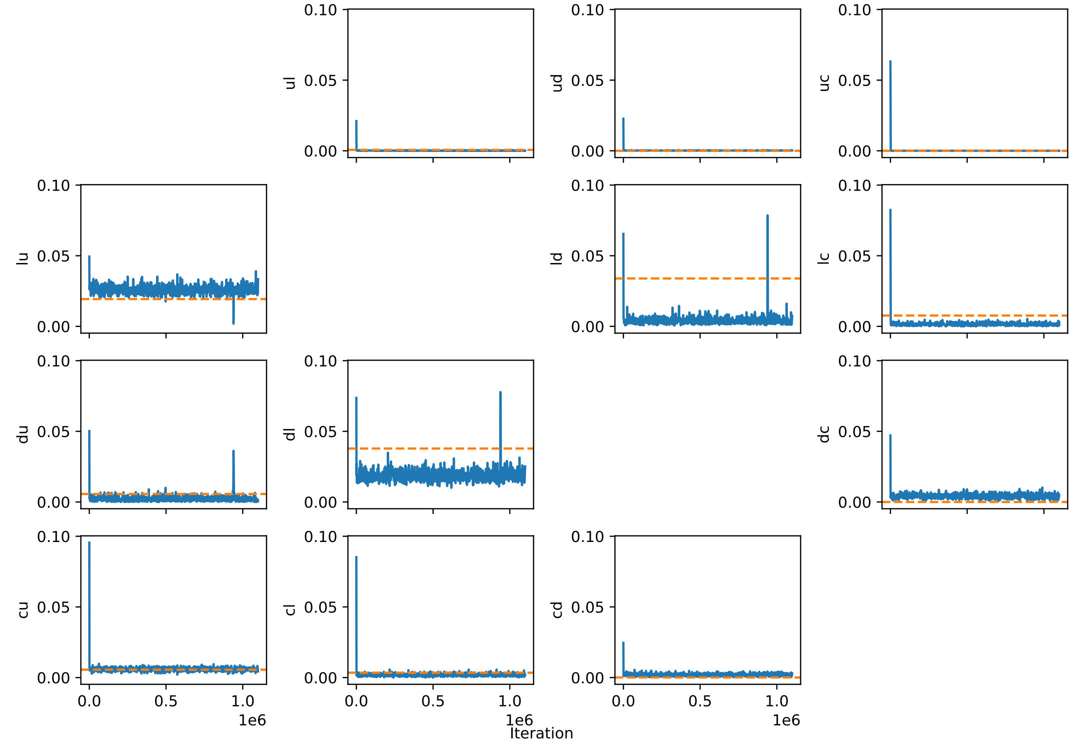


FIG. S19. **Janssens *et al.* [8] species trace.** Uniform prior between 0 and 0.1. Total no. iterations, 1100000. Burn in period, 100000 iterations. Sample period 1000 iterations. ML estimates given by the orange dashed line. Calculated using BayesTraits V4.1.2 [11].

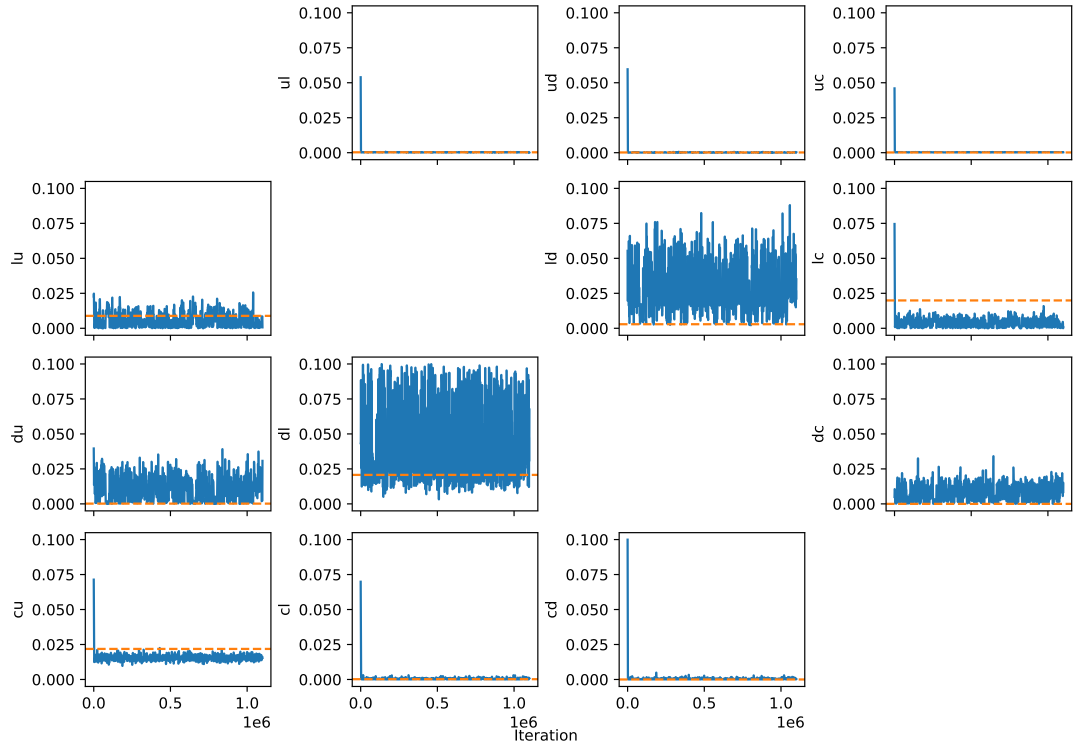


FIG. S20. **Janssens *et al.* [8] genus trace.** Uniform prior between 0 and 0.1. Total no. iterations, 1100000. Burn in period, 100000 iterations. Sample period 1000 iterations. ML estimates given by the orange dashed line. Calculated using BayesTraits V4.1.2 [11].

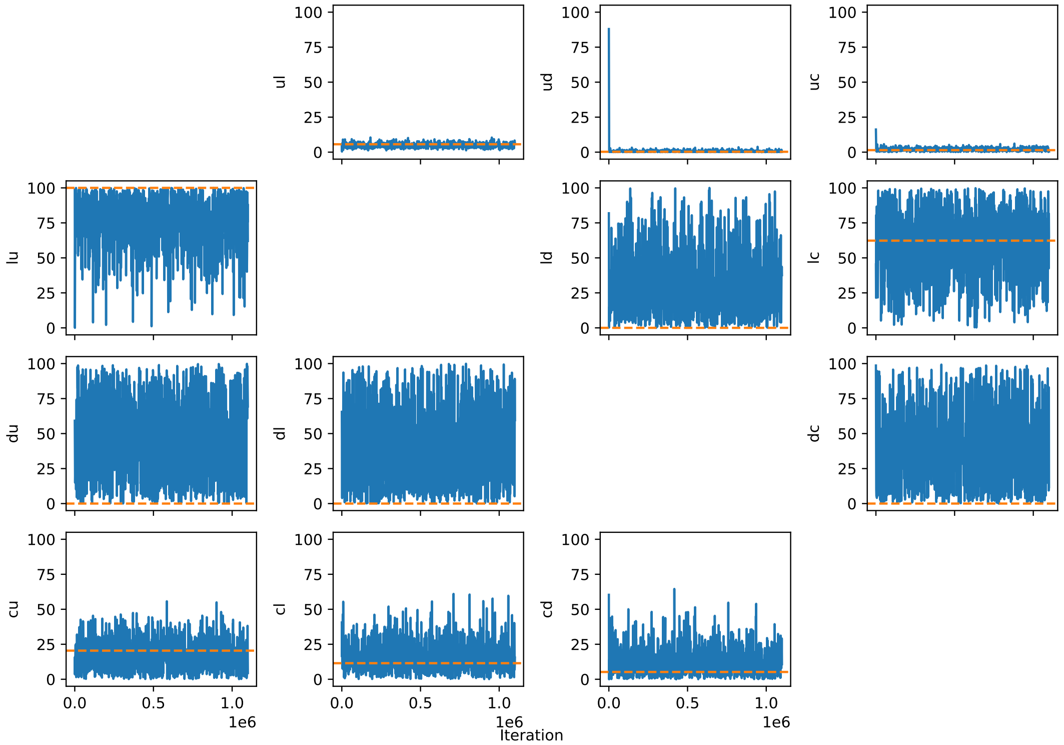


FIG. S21. **Geeta *et al.* [6] trace.** Uniform prior between 0 and 100. Total no. iterations, 1100000. Burn in period, 100000 iterations. Sample period 1000 iterations. ML estimates given by the orange dashed line. Calculated using BayesTraits V4.1.2 [11].

#### B. PCA occupancy for higher dimensions

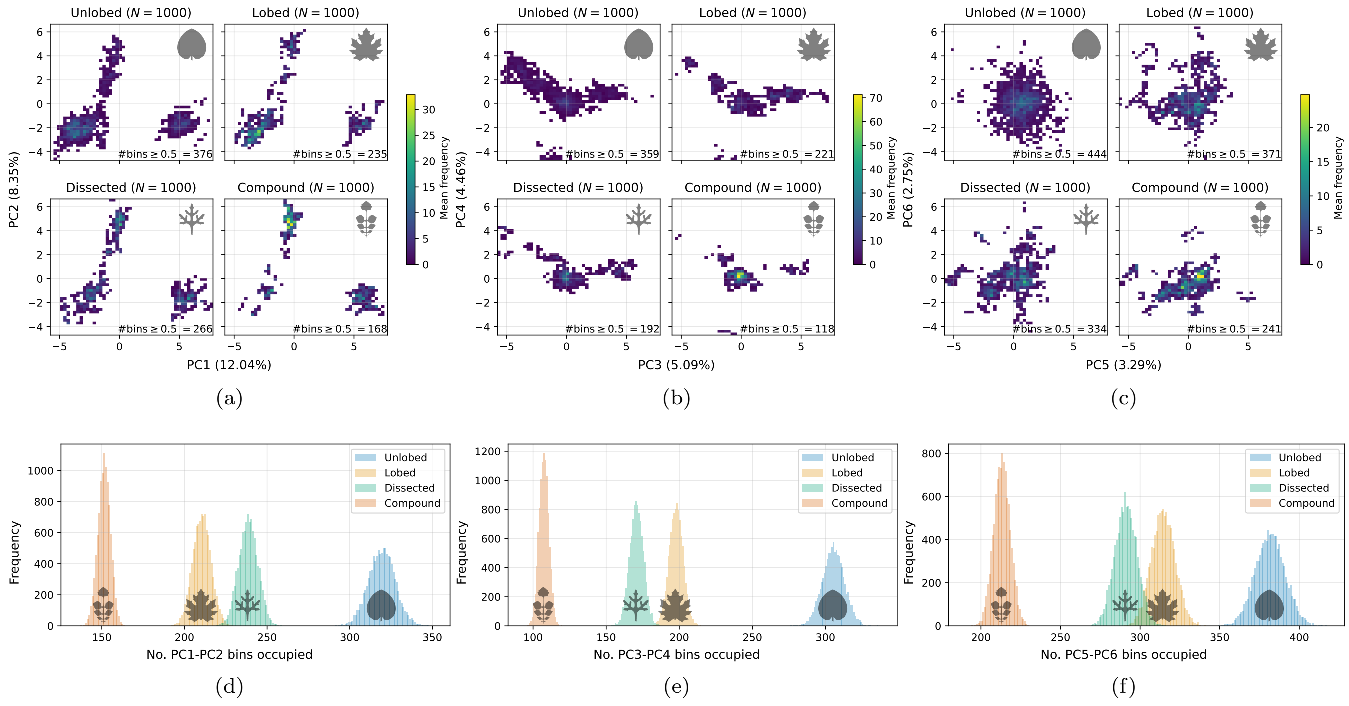


FIG. S22. **The unlobed category occupies a larger volume within parameter space along principal components 1-6.** Furthermore, the regions occupied by more complex shapes continue to overlap substantially with the unlobed domain at higher dimensions.

##### C. Detailed balance analysis

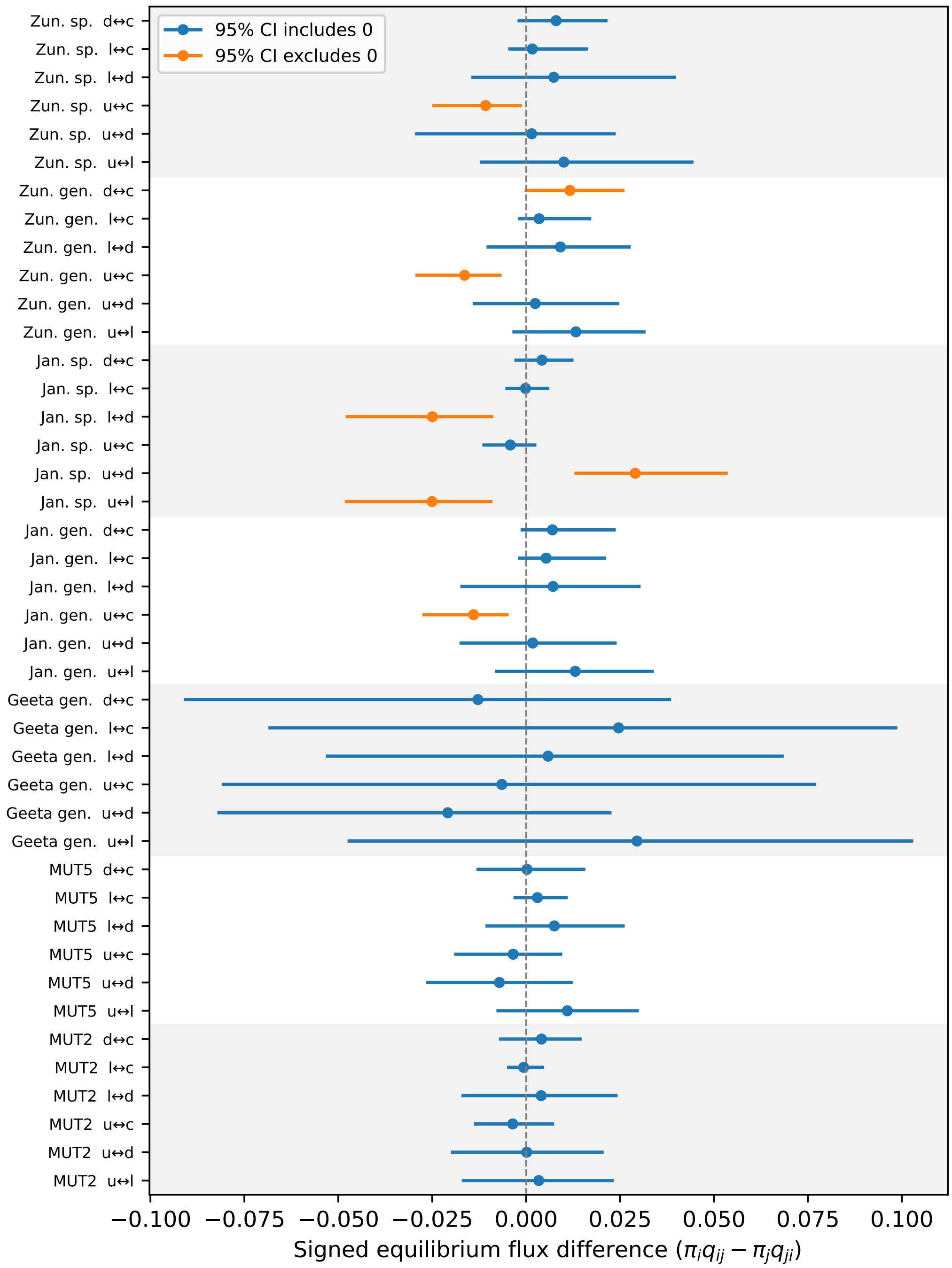


FIG. S23. **Posterior signed equilibrium flux difference for rate matrices inferred from all simulated and phylogenetic datasets.** Each point is the posterior median of $\pi_i q_{ij} - \pi_j q_{ji}$ for a pair of states, where $\boldsymbol{\pi}$ is the stationary distribution over the four states and $\boldsymbol{Q}$ is the posterior rate matrix. The row vector $\boldsymbol{\pi}$ is defined for each posterior draw of $\boldsymbol{Q}$ by solving $\boldsymbol{\pi Q} = 0$ where $\sum_i \pi_i = 1$. Horizontal bars show 95% credible intervals. Colour shows whether the interval includes zero, indicating consistency with detailed balance, or excludes zero, indicating imbalance. Alternating background bands group state pairs from the same dataset. Simulation transition rates are consistent with detailed balance, as expected given the symmetric mutational schemes (algs. 1 and S2). Some phylogenetic state pairs are inconsistent with detailed balance, implying potential non-reversibility in the inferred process; however, the affected pairs differ between phylogenetic trees. A positive flux difference means that flux is larger from the first to the second state.

### S4. A SHORT LITERATURE REVIEW OF THE ADAPTIVE SIGNIFICANCE OF LEAF SHAPE

#### A. Introduction

As the primary sites of photosynthesis, leaves are critical to plant fitness. They also display extraordinary variation in shape. It is therefore natural to ask to what extent this variation is the product of natural selection. Functional properties of leaves where shape may be relevant include thermoregulation, hydraulic efficiency, biomechanical stability, herbivory defence, light interception and nutrient cycling. Yet, despite an extensive and sophisticated literature, the extent to which leaf shape determines fitness, and hence reflects adaptation, remains the subject of considerable debate. In particular, a recurring difficulty in this literature is distinguishing evidence that morphology affects leaf function from evidence that the resulting functional differences affect fitness, and ultimately from evidence that selection on those differences explains evolutionary patterns of leaf shape.

In this short overview we briefly survey the literature on the role of adaptation in shaping three different classes of shape variables. The first is leaf size, which can vary enormously, from less than $1\,\mathrm{mm}^2$ to over $1\,\mathrm{m}^2$, and correlates quite strongly with climate: it was noted more than a century ago that leaves tend to be large in hot, sunny and wet climates, and small in hot, sunny and arid climates. The second class is marginal leaf teeth, which are small oscillatory patterns or serrations on the margin of leaves. The presence and number of teeth are thought to correlate with mean annual temperature, and have historically been used as a proxy to infer paleoclimate trends. The main class we treat here comprises dissection traits, which are captured by our unlobed, lobed, dissected and compound categories. We note that each of these four dissection categories can exhibit large variations in size, and can be toothed or untoothed.

#### B. Size

Leaf size is thought to be particularly important for thermoregulation, hydraulic performance, and light capture. In the case of thermoregulation, leaves need to regulate their temperature to maintain efficient photosynthesis and respiration, prevent damage from heat stress and freezing, and balance heat loss against water loss through transpiration [12–15]. Heat exchange between a leaf and the surrounding air depends partly on the thickness of the boundary layer of relatively still air at the leaf surface [16]. Smaller leaves generally have thinner boundary layers and therefore exchange heat with the surrounding air more rapidly than larger leaves. A recent large-scale study which analysed 7,670 species from 682 sites worldwide [14] argued that the leaf-to-air temperature difference plays a key role in explaining why large leaves predominate in warm, wet environments, whereas small leaves are common in hot, arid environments and at high altitudes. Interestingly, although these authors did separate out simple from compound leaf shapes, both dissection categories showed similar trends with climate variables.

Leaf size may also influence hydraulic performance. For example, Scoffoni *et al.* [17] used simulations and comparisons among 10 species with different drought tolerance to show that smaller leaves with greater major-vein density showed reduced hydraulic vulnerability and greater tolerance to vein embolism. Thus, variation in size may influence both the thermal and hydraulic conditions under which leaves can function. See also Sack and Scoffoni [18].

Size is also associated with trade-offs in light capture and resource allocation. Increasing leaf area can increase total photosynthetic surface [19], but leaf mass increases with area while proportional gains in area diminish, implying increasing construction costs for larger leaves [20]. Leaf size additionally interacts with canopy architecture. Smaller leaves may be associated with shorter petioles and greater crowding around stems, potentially increasing self-shading [21]. The leaf economic spectrum further demonstrates that investment in leaf structure is associated with major differences in leaf lifespan and photosynthetic return [22].

In conclusion, leaf size can have substantial functional consequences and is likely shaped by trade-offs between heat exchange, hydraulic efficiency, construction costs, and light capture. The evidence linking leaf size to ecologically important functions and environmental gradients is arguably stronger than the corresponding evidence for many aspects of leaf shape. Many of the functions affected by leaf size can also be modified by other aspects of leaf shape. This makes disentangling adaptive from non-adaptive variation more complex.

#### C. Teeth

Marginal teeth are shallow projections or serrations separated by sinuses, incised less than 25% of the distance to the midvein [23]. They are a much less pronounced feature than our dissection categories. Unlobed, lobed, dissected leaves, as well as the leaflets of compound leaves, can be either toothed or untoothed. Indeed, a close look at the first two unlobed leaves in fig. S6 reveals a toothed pattern in one and an untoothed pattern in the other, illustrating that the Runions model [1] can reproduce this feature on top of the dissection traits.

The presence or absence of leaf teeth has been of particular interest for over a century, due to an observed negative correlation with mean annual temperature (MAT), in particular for woody dicot species [24]. The association between climate and leaf tooth presence has been measured repeatedly [24–26], including by a study across 3,549 species and 6 continents [27]. The apparent simplicity and strength of this relationship led to its widespread use to estimate paleoclimate MAT. However, as independent methods for reconstructing past temperatures such as oxygen-isotope measurements and phytoplankton distributions were refined and recalibrated, discrepancies began to emerge between estimates of MAT based on fossil leaf margins and those obtained from these independent paleoclimate proxies. This motivated work to better understand and refine the relationship [28]. Multivariate approaches, particularly Climate-Leaf Analysis Multivariate Program (CLAMP) [29] or digital leaf physiognomy [26], were subsequently developed and refined to incorporate additional aspects of leaf morphology [26, 29]. New features, including the shape factor ($4\pi \times$blade area/perimeter$^2$), a measure of roundness, were found to correlate with MAT, albeit with a modest amount of variance explained ($r^2 = 0.22, P < 0.001$) [26].

However, there are several complicating factors. An important study of 569 species-site pairs that explicitly took phylogenetic models into account demonstrated that ancestry of plant species also plays a significant role in all 21 leaf traits measured, including the abundance of teeth [30]. Accounting for shared ancestry decreased the slope of the regression between the presence of teeth and mean annual temperature by an order of magnitude (in log space) to almost a flat line with wider confidence intervals. This suggests that the geographical correlation contains a substantial phylogenetic component, although it does not by itself distinguish whether this arises from phylogenetic constraint, climatic sorting of lineages, or repeated adaptive convergent evolution of toothed or untoothed margins. Subsequent studies suggest that an association between temperature and tooth presence remains after accounting for phylogeny, potentially mediated by venation architecture [31], although the picture remains complex [32], and clear counterexamples to the MAT correlation with teeth can also be found in certain species [28, 33].

Interestingly, Little *et al.* [30] also found that relationships between MAT and more gross shape variables such as shape factor changed little after phylogenetic correction, making these associations more consistent with repeated environmental responses across lineages. However, these correlations are not that strong in the first place. Moreover, the shape factor captures roundness rather than the specific distinctions among unlobed, lobed, dissected and compound leaves we consider in this study. Overall, an emerging consensus in the field is that evolutionary history and climatic selection jointly contribute to the observed relationship rather than that the relationship is simply a phylogenetic artefact. Modern work on paleoenvironmental reconstruction combines different leaf shape traits with phylogenetic, biogeographic and other analyses rather than relying on the fraction of leaves with teeth alone.

Despite the complications above, many functional hypotheses have been proposed to explain the climate association. Firstly, enhanced early-season photosynthesis at leaf teeth could be particularly advantageous in climates with short growing seasons [34]. Secondly, the ‘support-supply’ hypothesis posits that cooler climates with short growing seasons favour thinner leaves with stronger veins that penetrate to the margin for which the optimal shape of the surrounding tissue is wedge-like [35]. Thirdly, the ‘bud-packing’ hypothesis postulates that cold climates favour dense packing of leaf primordia in buds which results in more teeth due to physical forces from folding [36]. However, the evidence for these is mixed. Although Royer and Wilf [34] show enhanced photosynthesis and transpiration in toothed margins, early in the growing season, the extent to which this increases fitness is unclear. Moreover, Zohner *et al.* [31] found that leaf thickness does not predict venation and teeth, contradicting the support-supply hypothesis. They also found that primordium number in buds does not predict margin state, contradicting the bud-packing hypothesis.

In summary, several explanations for the correlation between leaf teeth and MAT remain viable. Teeth may be direct targets of selection because of some functional property they confer, but the relationship could also arise from a more complex combination of adaptive effects, including effects that partly cancel one another, or from relatively weak selection. Alternatively, teeth may be exaptations whose association with climate reflects selection on other aspects of leaf function. Distinguishing among these possibilities remains difficult. Thus, even for leaf toothiness, where the geographical association with climate is unusually strong and well replicated, identifying the selective mechanism responsible for that association remains difficult. This provides a useful point of comparison for the more pronounced dissection traits considered below, for which geographical correlations are generally less well established.

#### D. Dissection and compoundness

In this section we look in more detail at the literature linking leaf dissection traits, measured in our case through the lobed, unlobed, dissected and compound leaf shapes, to a series of different functions of leaves.

##### 1. Thermoregulation

Thermoregulation is frequently proposed as an important driver of leaf-shape evolution [37]. In particular, complex leaves have been hypothesised to behave aerodynamically like many small leaves, resulting in faster heat transfer to the environment [38]. Laboratory studies using artificial leaves have suggested that leaf heat exchange with the environment is strongly influenced by shape [16]. For example, Vogel [16] found that lobed copper plates were more effective heat dissipators than circles. However, this approach ignores other important traits of real leaves (e.g. water content, absorptance) that have large effects on the actual operating temperatures of leaves in natural environments [37].

A more recent study by Leigh *et al.* [38] measured multiple temperature variables in field conditions for 68 Proteaceae species, finding that effective leaf width was a strong predictor of leaf thermal dynamics, whereas leaf margin complexity had no or weak effects. Thus, although dissection can affect convective heat transfer under controlled conditions, its contribution to leaf temperature in vivo may be small relative to other morphological and physiological traits.

Ivyleaf morning glory (*Ipomoea hederacea*) provides an unusually well-studied test case. An investigation of lobed and unlobed genotypes of Ivyleaf morning glory (*Ipomoea hederacea*) [39] revealed that while both shapes maintain similar temperatures, unlobed phenotypes had significantly increased stomatal conductance in warmer, sunnier conditions. This suggests that unlobed leaves may rely more on latent cooling via transpiration for temperature regulation compared to lobed leaves. The fact that lobed leaves maintain the same temperature without increased stomatal conductance is consistent with a different cooling strategy, perhaps enhanced convective cooling facilitated by the lobed shape, to conserve water. This highlights how thermoregulation depends on a range of traits and even if leaf shape is one of them, suboptimal leaf shapes can arise due to trade-offs with these other traits.

*Ipomoea hederacea* also provides an instructive example of the difficulty of inferring selection from geographical patterns. The species has a clear latitudinal cline in North America, from lobed genotypes in the north to heart-shaped unlobed genotypes in the south, that is known to be determined by a single locus [40]. Campitelli and Stinchcombe [40] showed that the leaf shape locus displays clinal variation and neutral loci do not, consistent with the leaf shape cline being caused by selection on leaf shape. However, it is possible that selection is acting instead on a different locus in linkage disequilibrium with the leaf shape locus. They suggest that the agent of selection could be temperature. However, contrary to previous studies that predict that lobes are favoured in hot environments, the lobed genotypes are found more frequently in the colder north end of the cline. To identify the selective agent for this cline, Campitelli and Stinchcombe [41] planted 1,680 lobed-unlobed F3 hybrids with varying leaf shape in the north and measured their fitness as well as other selective agents including temperature and herbivory. They found that none of the potential selective agents affected the different leaf shape genotypes differently. This could be because the true selective agent was not among those investigated, or that the cline is not due to selection on leaf shape but is perhaps instead due to selection on another trait in linkage disequilibrium. This study highlights the difficulty in establishing selection as a cause, despite correlations between leaf shape and environmental variables.

##### 2. Hydraulic efficiency

Leaf shape has also been proposed to influence hydraulic efficiency. Photosynthetic capacity is directly impacted by the efficiency of water transport through leaves, which alone accounts for $\geq 30\%$ of the total resistance to water flow through plants [42]. Leaf structure, particularly venation architecture, strongly affects the strength of this resistance.

Sisó *et al.* [43] found that in 7 species of oak (*Quercus*), leaves with increased lobation had lower hydraulic resistance. It is thought that this is caused by a lower ratio of mesophyll tissue to vasculature in lobed leaves. This is consistent with the hypothesis that deeply lobed leaves may be a way of improving water balance in arid environments.

On the other hand, Sack and Frole [42] investigated the relationship between leaf structure and hydraulic resistance in 10 coexisting tropical rain forest tree species in Panama. They found that while resistance was negatively associated with vein density, there was no significant association with shape. Moreover, Scoffoni *et al.* [17] found, through simulations and comparisons among 10 diverse species with highly varying drought tolerance, that smaller leaves with higher major vein density had reduced hydraulic vulnerability and higher tolerance to vein embolism (the blocking of veins by air bubbles).

Therefore, while shape may have some effect on hydraulic efficiency, properties of the vasculature, such as vein density, may more directly determine hydraulic function.

##### 3. *Biomechanical stability*

Leaves are constantly subjected to various mechanical stresses from their environment, including static loads such as their own weight, accumulated snow or ice, and dynamic loads primarily from wind and rain. In order to maintain optimal fitness-critical functions such as photosynthesis, they need to withstand these stresses with minimal damage, and maintain optimal positioning for light interception. Much of the biomechanical literature concerns dimensions or petiole properties rather than dissection per se. However, it is possible that leaf shape affects the ability to withstand mechanical stress and therefore has been optimised by natural selection for mechanical robustness, particularly in high-stress environments (e.g. windy environments).

Tadrist and Darbois-Texier [44] found that they could predict scaling relationships among palm leaf length, width, thickness and petiole diameter with a mathematical model that maximises surface area under gravitational self-support constraints. This suggests that for this particular group of large-leaved plants, leaf dimensions may be constrained by optimisation for self-support. It is not clear how important other shape features, such as margin complexity, are for self-support.

Moreover, Louf *et al.* [45] predicted a relationship between blade aspect ratio and the relative bending and twisting rigidity of petioles, which was broadly consistent with measurements from four broad-leaved species. This is consistent with adaptive trade-offs between leaf shape and petiole mechanical properties as a result of mechanical stress from wind.

Niklas [46] found that leaves from wind-exposed and protected sugar maple trees (*Acer saccharum*) differed significantly in some aspects of shape. Leaves from wind-exposed trees had smaller surface areas and shorter and narrower petioles that were less rigid than petioles from leaves on protected trees. These differences are thought to be due to adaptive phenotypic plasticity in response to chronic mechanical perturbation by wind. The shorter, more flexible petioles allow for easier twisting and bending in response to the wind, reducing drag forces and the chance of mechanical tissue damage [47].

Different leaf shapes experience different levels of drag, relative to their surface area, in response to wind. Through wind-tunnel experiments, Vogel [48] found that pinnately compound leaves had lower drag than clusters of simple leaves by reconfiguring into a streamlined cylinder shape with alternately layered leaflets. This demonstrates a physical effect of leaf shape on drag and reconfiguration in wind, although whether this translates into differences in fitness remains to be established.

##### 4. *Herbivory defence*

Leaf herbivory is costly to plant fitness by reducing photosynthetic capacity. Leaves have evolved many features that deter herbivores such as chemical toxicity and physical barriers such as the waxy cuticle, thorns and trichomes. The role of leaf shape is less obvious; nevertheless, there is some evidence that it can impact herbivory.

For example, Higuchi and Kawakita [49] showed that the leaf-rolling weevil *Apoderus praecellens* could be deterred from cutting the leaves of its host genus *Isodon* by replacing leaves from an unlobed species (*I. trichocarpus*) typical of the genus with lobed leaves from a closely related species (*I. umbrosus*). They controlled for potential confounding differences between the plant species by cutting lobes into *I. trichocarpus* leaves to resemble *I. umbrosus*, and still found a preference for the non-lobed *I. trichocarpus* leaves among the female weevils.

Moreover, there is observational [50] and direct experimental [51] evidence that ovipositing butterflies use vision to recognise plant hosts by leaf shape, and can have a preference for some shapes over others. This shows that butterflies have the potential to exert selection on leaf shape in host species.

On the other hand, Campitelli *et al.* [52] investigated how alternative leaf morphs of the Ivyleaf morning glory *Ipomoea hederacea* affect the performance (e.g. insect biomass, leaf area consumed, digestive efficiency) of three different generalist insect herbivores. They found that the effect of shape on performance was highly variable between insect species, plant developmental stage and growth conditions. Therefore, while some insect herbivores may be able to consume certain leaf shapes more easily, it seems there is no consistent trend and other plant traits may play a more significant role.

##### 5. *Light interception and optimisation*

Leaves function as the primary organs for light harvesting in vascular plants, and are therefore of utmost importance to photosynthetic capacity. While the effects of several non-shape traits on photosynthetic capacity are well understood, the effect of shape per se is more uncertain [37].

Simulations of maize (*Zea mays*) have found that variation in leaf shape can alter light interception by up to 7% [53]. *Z. mays* leaves do not vary as substantially in shape as those of other lineages, suggesting that the effect on light interception may be greater for other plants with more shape variation.

Further simulations in oil palm found that parameters related to leaf area had the largest effect on light interception and carbon assimilation, and the effect of other shape parameters was less important [54].

Several studies on poplars have shown that some aspects of leaf shape can evolve to reduce self-shading within canopies. Aspen (*Populus tremuloides*) has evolved a flattened, non-rigid petiole oriented perpendicular to the leaf blade, resulting in enhanced leaf flutter even in light winds [55]. By directly measuring light intensity at different points within the canopy of two poplar species, Roden and Pearcy [56] showed that leaf flutter increased light penetration into the lower canopy, creating a more spatially even understory light environment. This can enhance whole canopy carbon gain, as understory poplar leaves are very efficient at utilising rapidly fluctuating light environments for photosynthesis [57]. Furthermore, evidence from simulations suggests that fluttering leaves at the top of the canopy have more uniform light interception than static leaves and therefore also potentially enhanced carbon gain [55]. This highlights how leaf shape can indirectly alter light interception by altering the mechanical response to wind. It is not clear how the shape of parts of the leaf besides the petiole could affect flutter.

Donoghue *et al.* [58] identified a link between leaf morphology and light levels in geographically isolated populations of *Viburnum* in Mexican and South American cloud forests. These populations showed convergent evolution of different leaf ecomorphs, consistent with repeated adaptive evolution into the same niches across these isolated locations. However, the authors have not ruled out neutral evolution as a potential cause. By measuring environmental variables in these isolated populations, they identified a positive correlation between large leaves, lower light levels and higher wetness. This is consistent with the hypothesis that larger leaves are an adaptation to lower light environments, but does not prove any differences in fitness associated with leaf shape. Moreover, the main correlation they found was related to size rather than shape.

##### *6. Nutrient cycling*

Deciduous trees shed their leaves annually, avoiding the cost of maintaining foliage under the stress of winter. However, this results in these trees losing 40% of their assimilated carbon and substantial amounts of nitrogen, phosphorus, potassium and other nutrients [59]. Given the potential nutritional value of these leaves, it is possible that there is a fitness benefit to trees that deposit their leaves nearby, such that their nutrients can be recycled and local soil conditions are improved. Biviano and Jensen [59] investigated the hypothesis that symmetric and unlobed leaves may have evolved as a strategy to increase leaf settling speed such that fallen leaves land closer to the tree of origin, facilitating nutrient retention. Through experiments with paper leaves, they found that highly symmetric shapes settled the fastest, with increasing lobedness decreasing settling speed. Asymmetric shapes settled substantially more slowly and were largely insensitive to changes in lobedness. This is consistent with the notion that leaf shape may also be under selection to facilitate nutrient recycling by maximising settling speed.

##### *7. Conclusion*

While the evidence for adaptation is much weaker for these dissection categories than it is for leaf size or even teeth, there remains substantial evidence that leaf dissection and compoundness can influence thermoregulation, hydraulics, mechanical behaviour, herbivore interactions and light interception. However, most studies establish functional effects rather than fitness consequences, and many are restricted to particular species or lineages. Moreover, in several cases leaf size, venation or petiole architecture appear to have stronger functional effects than dissection itself. Dissection may therefore be adaptive in particular ecological contexts, and different selective pressures may favour similar shapes in different contexts, but there is currently limited evidence for a general selective explanation for its broad evolutionary distribution. Absence of evidence is not necessarily evidence of absence. So, while there may still be an adaptive explanation, or several explanations, for the higher evolutionary transition rates towards unlobed shapes than complex shapes—lobed, dissected and compound—the current literature does not offer a clear answer to what this might be.

---

[1] A. Runions, M. Tsiantis, and P. Prusinkiewicz, A common developmental program can produce diverse leaf shapes, New Phytologist **216**, 401 (2017).

[2] M. Pagel, Detecting correlated evolution on phylogenies: A general method for the comparative analysis of discrete characters, Proc. R. Soc. Lond. B **255**, 37 (1994).
[3] J. Felsenstein, Evolutionary trees from DNA sequences: A maximum likelihood approach, Journal of Molecular Evolution **17**, 368 (1981).
[4] A. Meade and M. Pagel, Bayestraits manual (2024).
[5] M. Pagel, A. Meade, and D. Barker, Bayesian estimation of ancestral character states on phylogenies, Systematic Biology **53**, 673 (2004).
[6] R. Geeta, L. M. Dávalos, A. Levy, L. Bohs, M. Lavin, K. Mummenhoff, N. Sinha, and M. F. Wojciechowski, Keeping it simple: Flowering plants tend to retain, and revert to, simple leaves, New Phytologist **193**, 481 (2012).
[7] A. R. Zuntini, T. Carruthers, O. Maurin, P. C. Bailey, K. Leempoel, G. E. Brewer, N. Epitawalage, E. Françoso, B. Gallego-Paramo, C. McGinnie, *et al.*, Phylogenomics and the rise of the angiosperms, Nature **629**, 843 (2024).
[8] S. B. Janssens, T. L. Couvreur, A. Mertens, G. Dauby, L.-P. M. Dagallier, S. Vanden Abeele, F. Vandelook, M. Mascarello, H. Beeckman, M. Sosef, *et al.*, A large-scale species level dated angiosperm phylogeny for evolutionary and ecological analyses, BDJ **8**, e39677 (2020).
[9] Naturalis Biodiversity Center, Naturalis Bioportal (2024).
[10] D. Foreman-Mackey, D. W. Hogg, D. Lang, and J. Goodman, Emcee: The MCMC hammer, Publications of the Astronomical Society of the Pacific **125**, 306 (2013).
[11] M. Pagel, C. O'Donovan, and A. Meade, General statistical model shows that macroevolutionary patterns and processes are consistent with Darwinian gradualism, Nat Commun **13**, 1113 (2022).
[12] J. Berry and O. Bjorkman, Photosynthetic response and adaptation to temperature in higher plants, Annu. Rev. Plant. Physiol. **31**, 491 (1980).
[13] T. J. Givnish, Comparative studies of leaf form: Assessing the relative roles of selective pressures and phylogenetic constraints, New Phytologist **106**, 131 (1987).
[14] I. J. Wright, N. Dong, V. Maire, I. C. Prentice, M. Westoby, S. Díaz, R. V. Gallagher, B. F. Jacobs, R. Kooyman, E. A. Law, M. R. Leishman, Ü. Niinemets, P. B. Reich, L. Sack, R. Villar, H. Wang, and P. Wilf, Global climatic drivers of leaf size, Science **357**, 917 (2017).
[15] T. M. Perez and K. J. Feeley, Photosynthetic heat tolerances and extreme leaf temperatures, Functional Ecology **34**, 2236 (2020).
[16] S. Vogel, Convective cooling at low airspeeds and the shapes of broad leaves, J Exp Bot **21**, 91 (1970).
[17] C. Scoffoni, M. Rawls, A. McKown, H. Cochard, and L. Sack, Decline of leaf hydraulic conductance with dehydration: Relationship to leaf size and venation architecture, Plant Physiology **156**, 832 (2011).
[18] L. Sack and C. Scoffoni, Leaf venation: Structure, function, development, evolution, ecology and applications in the past, present and future, New Phytologist **198**, 983 (2013).
[19] D. J. Watson, The dependence of net assimilation rate on leaf-area index, Annals of Botany **22**, 37 (1958).
[20] K. J. Niklas, E. D. Cobb, Ü. Niinemets, P. B. Reich, A. Sellin, B. Shipley, and I. J. Wright, "diminishing returns" in the scaling of functional leaf traits across and within species groups, Proc. Natl. Acad. Sci. U.S.A. **104**, 8891 (2007).
[21] D. S. Falster and M. Westoby, Leaf size and angle vary widely across species: What consequences for light interception?, New Phytologist **158**, 509 (2003).
[22] I. J. Wright, P. B. Reich, M. Westoby, D. D. Ackerly, Z. Baruch, F. Bongers, J. Cavender-Bares, T. Chapin, J. H. Cornelissen, M. Diemer, *et al.*, The worldwide leaf economics spectrum, Nature **428**, 821 (2004).
[23] B. Ellis, D. Daly, L. Hickey, K. Johnson, J. Mitchell, P. Wilf, and S. Wing, *Manual of Leaf Architecture* (Cornell University Press CABI, GB, 2009).
[24] I. W. Bailey and E. W. Sinnott, The climatic distribution of certain types of angiosperm leaves, American Journal of Botany **3**, 24 (1916).
[25] D. L. Royer, P. Wilf, D. A. Janesko, E. A. Kowalski, and D. L. Dilcher, Correlations of climate and plant ecology to leaf size and shape: Potential proxies for the fossil record, American Journal of Botany **92**, 1141 (2005).
[26] D. J. Peppe, D. L. Royer, B. Cariglino, S. Y. Oliver, S. Newman, E. Leight, G. Enikolopov, M. Fernandez-Burgos, F. Herrera, J. M. Adams, E. Correa, E. D. Currano, J. M. Erickson, L. F. Hinojosa, J. W. Hoganson, A. Iglesias, C. A. Jaramillo, K. R. Johnson, G. J. Jordan, N. J. B. Kraft, E. C. Lovelock, C. H. Lusk, Ü. Niinemets, J. Peñuelas, G. Rapson, S. L. Wing, and I. J. Wright, Sensitivity of leaf size and shape to climate: Global patterns and paleoclimatic applications, New Phytologist **190**, 724 (2011).
[27] D. L. Royer, D. J. Peppe, E. A. Wheeler, and Ü. Niinemets, Roles of climate and functional traits in controlling toothed vs. untoothed leaf margins, American Journal of Botany **99**, 915 (2012).
[28] R. J. Burnham and G. S. Tonkovich, Climate, leaves, and the legacy of two giants, New Phytologist **190**, 514 (2011).
[29] J. A. Wolfe, *A Method of Obtaining Climatic Parameters from Leaf Assemblages*, Tech. Rep. 2040 (U.S. G.P.O. ; For sale by USGS Map Distribution,, 1993).
[30] S. A. Little, S. W. Kembel, and P. Wilf, Paleotemperature proxies from leaf fossils reinterpreted in light of evolutionary history, PloS One **5**, e15161 (2010).
[31] C. M. Zohner, E. Ramm, and S. S. Renner, Examining the support–supply and bud-packing hypotheses for the increase in toothed leaf margins in northern deciduous floras, American Journal of Botany **106**, 1404 (2019).
[32] Y. Li, Z. Wang, X. Xu, W. Han, Q. Wang, and D. Zou, Leaf margin analysis of Chinese woody plants and the constraints on its application to palaeoclimatic reconstruction, Global Ecology and Biogeography **25**, 1401 (2016).

[33] H. A. Frye, K. Mocko, T. E. Moore, C. D. Schlichting, and C. S. Jones, Leaf margins in a deciduous lineage from the Greater Cape Floristic Region track climate in unexpected directions, American Journal of Botany **107**, 735 (2020).
[34] D. L. Royer and P. Wilf, Why do toothed leaves correlate with cold climates? gas exchange at leaf margins provides new insights into a classic paleotemperature proxy, International Journal of Plant Sciences **167**, 11 (2006).
[35] T. J. Givnish and R. Kriebel, Causes of ecological gradients in leaf margin entirety: Evaluating the roles of biomechanics, hydraulics, vein geometry, and bud packing, American Journal of Botany **104**, 354 (2017).
[36] E. J. Edwards, E. L. Spriggs, D. S. Chatelet, and M. J. Donoghue, Unpacking a century-old mystery: Winter buds and the latitudinal gradient in leaf form, American Journal of Botany **103**, 975 (2016).
[37] A. B. Nicotra, A. Leigh, C. K. Boyce, C. S. Jones, K. J. Niklas, D. L. Royer, and H. Tsukaya, The evolution and functional significance of leaf shape in the angiosperms, Functional Plant Biol. **38**, 535 (2011).
[38] A. Leigh, S. Sevanto, J. Close, and A. Nicotra, The influence of leaf size and shape on leaf thermal dynamics: Does theory hold up under natural conditions?, Plant Cell & Environment **40**, 237 (2017).
[39] Y. K. Singhal, J. A. Boyle, and J. R. Stinchcombe, Differences in stomatal conductance between leaf shape genotypes of *Ipomoea hederacea* suggest divergent ecophysiological strategies (2025).
[40] B. E. Campitelli and J. R. Stinchcombe, Natural selection maintains a single-locus leaf shape cline in Ivyleaf morning glory, *Ipomoea hederacea*, Molecular Ecology **22**, 552 (2013).
[41] B. E. Campitelli and J. R. Stinchcombe, Testing potential selective agents acting on leaf shape in *Ipomoea hederacea*: Predictions based on an adaptive leaf shape cline, Ecology and Evolution **3**, 2409 (2013).
[42] L. Sack and K. Frole, Leaf structural diversity is related to hydraulic capacity in tropical rain forest trees, Ecology **87**, 483 (2006).
[43] S. Sisó, J. Camarero, and E. Gil-Pelegrín, Relationship between hydraulic resistance and leaf morphology in broadleaf *Quercus* species: A new interpretation of leaf lobation, Trees **15**, 341 (2001).
[44] L. Tadrist and B. Darbois-Texier, Are leaves optimally designed for self-support? an investigation on giant monocots, Journal of Theoretical Biology **396**, 125 (2016).
[45] J.-F. Louf, L. Nelson, H. Kang, P. N. Song, T. Zehnbauer, and S. Jung, How wind drives the correlation between leaf shape and mechanical properties, Sci Rep **8**, 16314 (2018).
[46] K. Niklas, Differences between *Acer saccharum* leaves from open and wind-protected sites, Annals of Botany **78**, 61 (1996).
[47] K. J. Niklas, A mechanical perspective on foliage leaf form and function, New Phytologist **143**, 19 (1999).
[48] S. Vogel, Drag and reconfiguration of broad leaves in high winds, J Exp Bot **40**, 941 (1989).
[49] Y. Higuchi and A. Kawakita, Leaf shape deters plant processing by an herbivorous weevil, Nat. Plants **5**, 959 (2019).
[50] M. D. Rausher, Search image for leaf shape in a butterfly, Science **200**, 1071 (1978).
[51] D. D. Dell'Aglio, M. E. Losada, and C. D. Jiggins, Butterfly learning and the diversification of plant leaf shape, Front. Ecol. Evol. **4**, 81 (2016).
[52] B. E. Campitelli, A. K. Simonsen, A. Rico Wolf, J. S. Manson, and J. R. Stinchcombe, Leaf shape variation and herbivore consumption and performance: A case study with *Ipomoea hederacea* and three generalists, Arthropod-Plant Interactions **2**, 9 (2008).
[53] D. Otto, S. Munz, E. Memic, J. Hartung, and S. Graeff-Hönninger, A computer vision approach for quantifying leaf shape of maize (*Zea mays* L.) and simulating its impact on light interception, Front. Plant Sci. **16**, 1521242 (2025).
[54] R. P. A. Perez, J. Dauzat, B. Pallas, J. Lamour, P. Verley, J.-P. Caliman, E. Costes, and R. Faivre, Designing oil palm architectural ideotypes for optimal light interception and carbon assimilation through a sensitivity analysis of leaf traits, Annals of Botany **121**, 909 (2018).
[55] J. S. Roden, Modeling the light interception and carbon gain of individual fluttering aspen (*Populus tremuloides* Michx) leaves, Trees **17**, 117 (2003).
[56] J. S. Roden and R. W. Pearcy, Effect of leaf flutter on the light environment of poplars, Oecologia **93**, 201 (1993).
[57] J. S. Roden and R. W. Pearcy, Photosynthetic gas exchange response of poplars to steady-state and dynamic light environments, Oecologia **93**, 208 (1993).
[58] M. J. Donoghue, D. A. R. Eaton, C. A. Maya-Lastra, M. J. Landis, P. W. Sweeney, M. E. Olson, N. I. Cacho, M. K. Moeglein, J. R. Gardner, N. M. Heaphy, M. Castorena, A. S. Rivas, W. L. Clement, and E. J. Edwards, Replicated radiation of a plant clade along a cloud forest archipelago, Nat Ecol Evol **6**, 1318 (2022).
[59] M. D. Biviano and K. H. Jensen, Settling aerodynamics is a driver of symmetry in deciduous tree leaves, J. R. Soc. Interface. **22**, 20240654 (2025).
